# Rediversification Following Ecotype Isolation Reveals Hidden Adaptive Potential

**DOI:** 10.1101/2023.05.03.539206

**Authors:** Joao A Ascensao, Jonas Denk, Kristen Lok, QinQin Yu, Kelly M Wetmore, Oskar Hallatschek

**Affiliations:** Department of Bioengineering, University of California Berkeley, Berkeley, CA, USA; Department of Physics, University of California Berkeley Berkeley, CA, USA; Department of Integrative Biology, University of California Berkeley, Berkeley, CA, USA; Department of Immunology and Infectious Diseases, Harvard T.H. Chan School of Public Health, Boston, Massachusetts, United States; Environmental Genomics and Systems Biology Division, Lawrence Berkeley National Laboratory, Berkeley, CA; Peter Debye Institute for Soft Matter Physics, Leipzig University, 04103 Leipzig, Germany

## Abstract

Microbial communities play a critical role in ecological processes, and their diversity is key to their functioning. However, little is known about if communities can regenerate ecological diversity following species removal or extinction, and how the rediversified communities would compare to the original ones. Here we show that simple two-ecotype communities from the *E. coli* Long Term Evolution Experiment (LTEE) consistently rediversified into two ecotypes following the isolation of one of the ecotypes, coexisting via negative frequency-dependent selection. Communities separated by more than 30,000 generations of evolutionary time rediversify in similar ways. The rediversified ecotype appears to share a number of growth traits with the ecotype it replaces. However, the rediversified community is also different compared to the original community in ways relevant to the mechanism of ecotype coexistence, for example in stationary phase response and survival. We found substantial variation in the transcriptional states between the two original ecotypes, whereas the differences within the rediversified community were comparatively smaller, but with unique patterns of differential expression. Our results suggest that evolution may leave room for alternative diversification processes even in a maximally reduced community of only two strains. We hypothesize that the presence of alternative evolutionary pathways may be even more pronounced in communities of many species, highlighting an important role for perturbations, such as species removal, in evolving ecological communities.

## Introduction

Ecological diversification is the process by which a population or community of organisms evolves to occupy different ecological niches or habitats in a given ecosystem^1^. This diversification can occur in various ways, such as the development of different physical or behavioral adaptations that allow individuals to exploit different resources or tolerate different environmental conditions^2–4^. The potential for ecological diversification within a community typically hinges on factors such as environmental conditions, existing biodiversity, and the ecological interactions among resident species^4–7^. Microbial communities have proven particularly useful for studying the interplay of evolutionary and ecological processes underlying diversification due to their manageable time scales for reproduction and evolution^8–16^. Diversification may be influenced by the availability of unoccupied niches, often referred to as “ecological opportunities”^5,17^, which can become scarce when most niches are already occupied due to high diversity levels. Alternatively, the resident community can create new niches, enabling the establishment of novel species, suggesting that “ diversity begets diversity”^18^. Cross-feeding exemplifies this latter scenario, where species release metabolites that can foster the emergence of new species by creating exploitable niches^13,15,19–21^. Ecological interactions within microbial communities have been shown to have negative^11,12^, positive^13,15^, and even mixed effects^22^ on a community’s ability to diversify.

However, the stable coexistence of a novel species and its ancestor is not guaranteed and may depend on various community properties, such as metabolic trade-offs^21,23^. In experimental settings, ecological differentiation of a diversified ecotype is often indicated when an ecotype’s fitness inversely correlates with its frequency, i.e. displaying negative frequency-dependent fitness effects. Stable coexistence between the diversified ecotype and its ancestor is implied if it can invade at small frequencies and cannot invade at large frequencies.

Even when species can stably coexist, it does not guarantee that they will coexist indefinitely or at all locations. Species can migrate to new territories, potentially without other community members, or some species within the community may spontaneously go extinct. In either case, the community becomes perturbed, losing one or more members and potentially leaving ecological niches unfilled. Theoretical models suggest that perturbed communities may respond with a combination of ecological and evolutionary changes^24–27^. These evolutionary changes may include both directional and diversifying selection^27^, with newly evolved variants either replacing existing community members or coexisting alongside them. However, it remains unclear which communities have the potential to rediversify. Recently diversified communities may be more likely to rediversify following ecotype isolation, as they have recently arisen from an ancestor that underwent ecological diversification. However, as a community coevolves, the potential for rediversification might diminish, but this may not necessarily always the case. When rediversification does occur after species removal, there are two possible scenarios: (i) the community eventually rediversifies and returns to a state similar to the original community before the disturbance, or (ii) the perturbed community rediversifies and forms a community that is qualitatively different from the original one.

Here, we investigate the aforementioned questions surrounding rediversification using a minimal microbial model community of only two, naturally diversified *E. coli* strains. Specifically, we employ two strains derived from the *E. coli* Long-Term Evolution Experiment (LTEE), which was started by Dr. Richard Lenski and has been running for over 30 years or more than 70,000 generations^28^. An initially isogenic strain of *E. coli* was split into 12 replicate populations and propagated through daily dilutions in glucose minimal media (DM25). At the outset of the LTEE around 6.5k generations, it was found that one lineage, ara-2, spontaneously diversified into two lineages–*S* and *L*–that coexist via negative frequency dependence^29^. The ecotypes were named for the sizes of their colonies on certain agar plates, either small (*S*) or large (*L*). The *S* and *L* lineages inhabit distinct temporal and metabolic niches in the LTEE environment. During exponential phase, L grows more quickly on glucose, while S specializes in stationary phase survival and utilizes acetate, a byproduct of overflow metabolism^30,31^. Since their diversification, the lineages have persisted and evolved over time, exhibiting genetic, transcriptional, and metabolic divergence^29–36^. The LTEE-derived communities are ideal for our plan to investigate the possibility and potential patterns of rediversification over evolutionary time. We can revive the *S*-*L* community at 6.5k generations to probe rediversification right after emergence of the community, and compare with rediversification at later stages of the evolution experiment.

We found that when we isolated the *S* ecotype under certain conditions, it would spontaneously rediversify, giving rise to a new big colony ecotype *S*_*B*_, even if we used *S* clones separated by more than 30,000 generations of evolutionary time. The new ecotype, *S*_*B*_, displays hallmarks of ecological differentiation, including negative frequency-dependent fitness effects when with its ancestral *S* clone. We dissected the new, rediversified community, and found that while *S*_*B*_ shares a number of traits with both *L* and *S*, it also behaves in entirely new ways. Our findings suggest that even in a maximally reduced community of only two strains, evolution may leave room for alternative diversification processes, suggesting a hidden adaptive potential only revealed by ecotype removal. This raises the possibility that perturbations, such as species removal, could play an important role in evolving ecological communities by creating opportunities for alternative evolutionary pathways.

## Results

### *S* can quickly diversify into a new ecotype

The ability of the *S* ecotype to emerge and coexist with the *L* ecotype in the LTEE has been attributed to its proficiency in scavenging acetate released from overflow metabolism during glycosis, as well as its ability to survive and thrive during stationary phase^30,37^. It has been proposed that the *L*-*S* and similar polymorphisms may arise because of a fundamental, hard-to-break trade-off between glucose and acetate growth rates in *E. coli*^21,38^. Based on these explanations, one may suspect, that after removing either *L* or *S* in the two species community, the community may eventually rediversify and will eventually approach a two-species community similar to the original *L*-*S* community.

We performed a simple experiment where we cultured an *S* clone isolated around 6.5k generations, immediately after the ara-2 lineage diversified into *S* and *L*, in glucose minimal media (DM25) for approximately 60 generations (9 days), with 12 biological replicates. To visualize colony morphologies of the resulting cultures, we plated the cultures on tetrazolium arabinose (TA) agar plates. Surprisingly, 2 of the independent cultures displayed a mixture of large and small colonies (Figure 1A).

**Figure 1.**
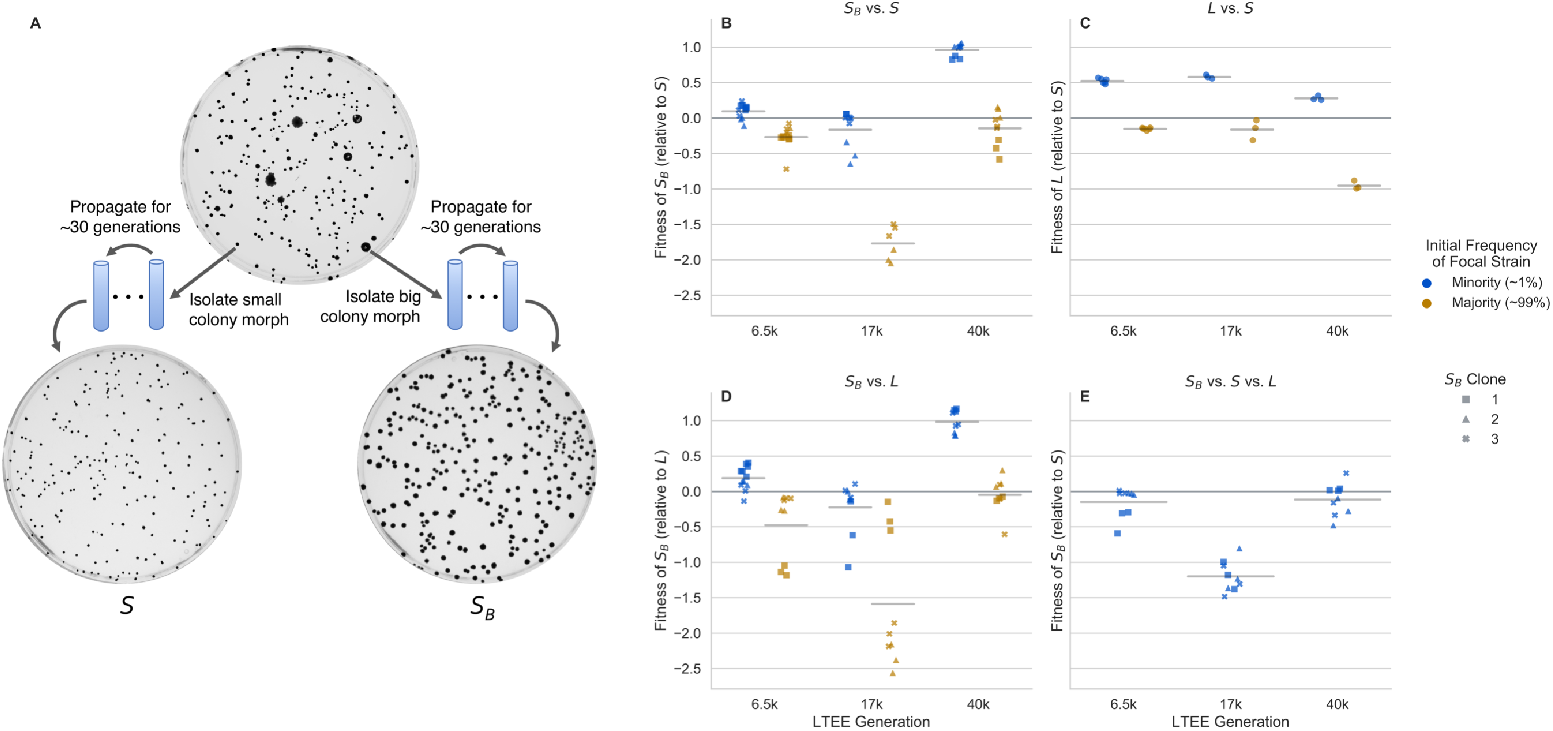
Emergence of the stably heritable *S*_*B*_ morph and frequency-dependent fitness effects. (**A**) Big colony morphs can arise in *S* cultures derived from 3 different LTEE timepoints, separated by more than 30,000 generations of evolution (6.5k clones are shown here as an example). When both small and big colonies are isolated and propagated in liquid DM25 culture for about 30 generations, then plated on TA agar plates, we see that the colony size is heritable. (**B-D**) Reciprocal invasion experiments, measuring relative fitness of clones when they are in the minority of the population (approximately 1%) or in the majority (approximately 99%). Each point represents a biological replicate, horizontal lines represent mean across all points. Competitions between (**B**) *S*_*B*_ clones and *S*, (**C**) *S* and *L*, and (**D**) *S*_*B*_ and *L*. We generally see negative frequency dependent fitness effects across all strains and competitions. (**E**) Triple competition between *S*_*B*_, *S*, and *L*, where *L*and *S*_*B*_ are near their equilibrium frequencies and *S*_*B*_ in the minority (around 1%). In this “full” community, *S*_*B*_ generally cannot invade, as most fitnesses are less than 0.

After eliminating contamination possibilities by sequencing several diagnostic genetic loci, we examined whether the large colony phenotype was heritable. We isolated several large and small colonies and propagated them in DM25 for around 30 generations (5 days). The phenotype appeared to be stably heritable for all selected colonies. To avoid prematurely associating the larger colony phenotype with the *L* type, we referred to the emerging type in our experiments as *S*_*B*_, due to its large (big) colonies and its ancestor *S*.

To gain insights into the robustness of the observed rediversification after isolation of *S* over evolutionary timescales, we isolated *S* from later generations, spanning more than 30,000 generations of evolution. We repeated the same experiment with *S* clones from 17k and 40k generations with 24 independent cultures each; however, we did not see any noticeable emergence of big colonies after 60 generations. It is unclear why we did not see any big colonies; one possible explanation may be that the rate at which *S* morphs transition to *S*_*B*_ morphs may be low enough that we would need to have many more replicate cultures to observe rediversification (as in the 6.5k *S* clones). We previously noticed that 6.5k *S*_*B*_ clones grew much better in LB liquid media compared to *S* clones (potentially accounting for their bigger colonies sizes on similar agar plates). Thus, we sought to see if we could enrich for the appearance of *S*_*B*_ by growing 6.5k, 17k, and 40k *S* clones in LB liquid culture. Under these growing conditions, we indeed saw that *S*_*B*_ colonies appeared rapidly, within 1-3 days, in nearly all of the independent *S* cultures across the three LTEE timepoints (Figure S5). We attributed this to the higher fitness of *S*_*B*_ in LB, relative to *S* (Figure S6). The new *S*_*B*_ clones were again stably heritable for at least 30 generations.

The big colony phenotype *S*_*B*_ bears at least a superficial resemblance to *L*, which begs the question: do *S*_*B*_ and *S* represent genuinely different ecotypes, occupying different ecological niches, with the potential to coexist with each other? To answer this question, we performed reciprocal invasion experiments, where we mixed *S* and *S*_*B*_ clones at high and low frequencies, and tracked how their frequencies change via flow cytometry (see Methods), to estimate their relative, frequency-dependent fitness effects (Figure 1B). While relative frequencies of LTEE strains are typically measured by colony counting, we found significant bias (Figure S2) in frequency measurements of *S*/*L* when measured via colony forming units (CFUs). In contrast, we see that flow cytometry provides unbiased frequency measurements (Figure S1). We thus chose to use flow cytometry for all further measurements instead of CFUs, owing to its minimal bias and reduced measurement noise (Figures S1 and S2B).

We found that all *S*_*B*_ clones had negative frequency-dependent fitness differences when in competition with their parental *S* clone, a hallmark of ecological differentiation. These data suggest that many of the *S*_*B*_ clones can coexist with *S*, because relative fitness is greater than 0 at low frequencies and less than 0 at high frequencies. However, it is not clear if this is the case for all of the isolated *S*_*B*_ clones, as some have a relative fitness near or less than 0 at low frequencies. This may be because the aforementioned *S*_*B*_ clones either genuinely do not coexist with *S*, or perhaps they coexist at a frequency around or lower than the one where we took the measurements.

The frequency-dependent fitness differences between *S*_*B*_ and *S* were similar in magnitude to the fitness differences between *L* and *S* (Figure 1C). We also competed *S*_*B*_ against *L*, and again found frequency-dependent fitness differences (Figure 1D). However, if at least some *S*_*B*_ clones can invade both *S* and *L* when rare, why has the *S*_*B*_ morph not appeared in the ara-2 population of the LTEE, where *L* and *S* have been coexisting and coevolving for tens of thousands of generations? We hypothesized that *S*_*B*_ could not invade an already “full” community, and could only have the chance to invade when one of the ecotypes is removed. Indeed, when we performed a triple competition experiment, with *L* and *S* near their equilibrium frequency and *S*_*B*_ in the minority, we found that *S*_*B*_ could not invade the community (Figure 1E).

While we have shown that *S*_*B*_ spontaneously emerges from a monoclonal population of *S* and occupies a distinct ecological niche, it is not yet clear how *S*_*B*_ compares to *S* and *L*. In particular, we want to understand if *S*_*B*_ simply fills the same niche that *L* had occupied before removal, making it somewhat functionally equivalent to *L*. In the following, we will show that while *S*_*B*_ resembles *L* in some of its growth properties, it also shows clear differences that are critical for its coexistence with *S*.

### Within-cycle growth dynamics of cocultures

To better understand how ecological differentiation arises in the *S*_*B*_-*S* and *L*-*S* systems, we measured the within-cycle growth dynamics of *S*_*B*_, *L*, and *S* in coculture with each other via flow cytometry. The LTEE environment is a seasonal one^30,39^–every 24 hours, cultures are transferred 1:100 into fresh glucose minimal media. The populations spend the first part of the day in exponential phase; the remaining time, more than 2/3 of the day, is spent transitioning out of exponential phase and in stationary phase. It has been previously shown that *L* and *S* occupy different temporal niches from one another, where *L* specializes on exponential growth on glucose, and *S* specializes on stationary phase survival and growth on acetate. Thus, it is natural to ask how temporal variations in growth are similar or different in the *S*_*B*_-*S* system.

To perform the experiments, we propagated *S, S*_*B*_, and *L* separately in monocultures for two days, before mixing *S* with *S*_*B*_ and *S* with *L*, both at high and low frequencies. We mixed strains with their partners from the same LTEE generation. For simplicity, we only used *S*_*B*_ clone 1 for all experiments and LTEE generations. We propagated the cocultures for one more cycle to allow the populations to physiologically adapt to the new environment. At the end of the 24 hour cycle, we took a flow cytometry measurement of the culture, then split the cultures into biological replicates and diluted the cocultures 1:100 into fresh media. Afterwards, we took flow cytometry measurements from the cocultures approximately every hour for about eight hours, then we took one last measurement at the end of the 24 hour cycle (Figures 2, S7). We chose this design because the fastest dynamics occur during and right after exponential phase–the first 8 hours–while dynamics in stationary phase are much slower. The cultures were grown in a 37°C shaking water bath. We corrected the cell counts measured in flow cytometry by the total dilution rate.

**Figure 2.**
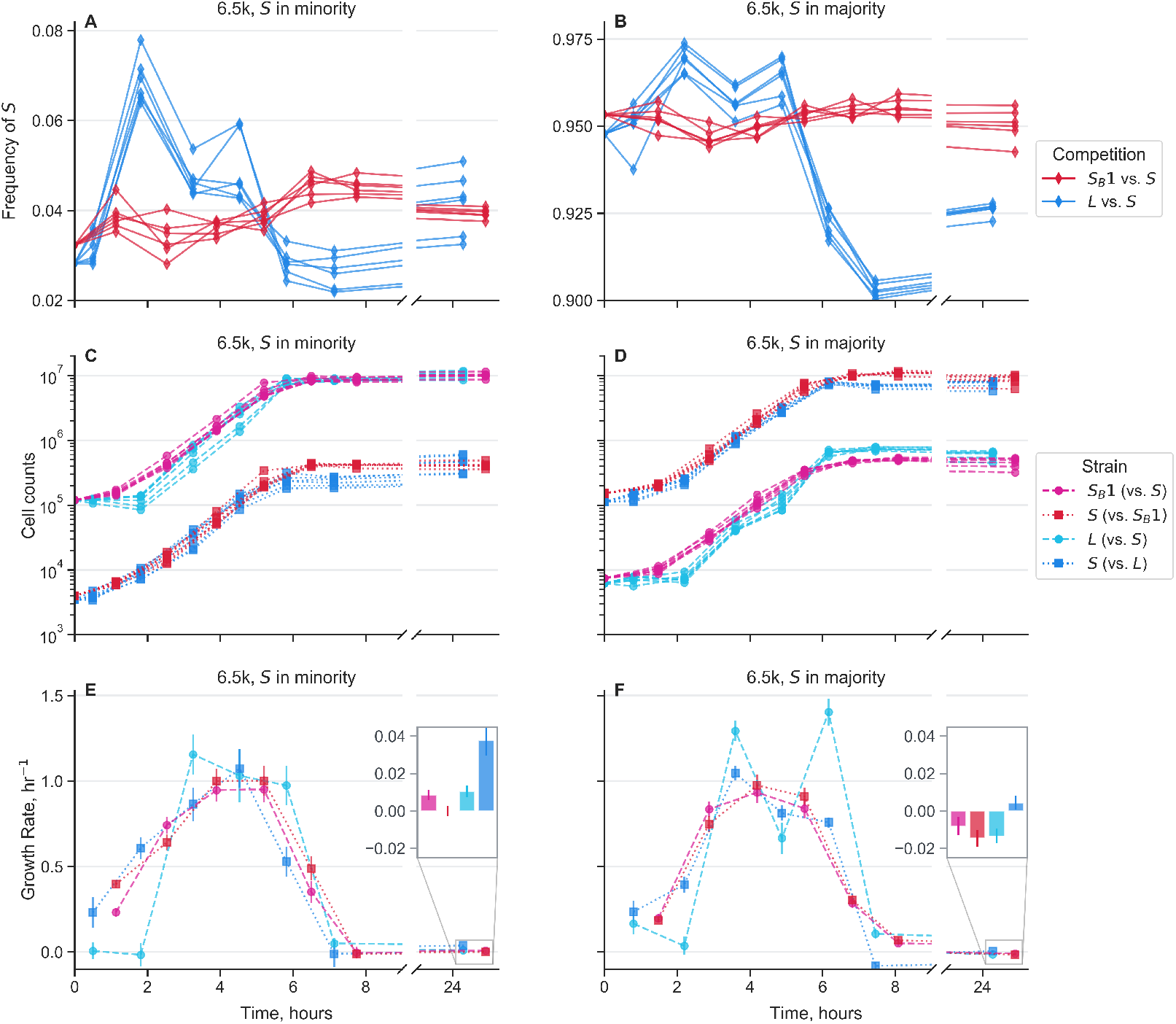
Growth dynamics of cocultures over the course of one twenty-four hour growth cycle. Measurements were taken approximately every hour via flow cytometry for the first eight hours after transfer into new media. An additional measurement was taken approximately 24 hours after the start of the cycle. Mixed *S*_*B*_ 1 with *S* along with *L* with *S*, all from 6.5k generations, where ecotypes were mixed both in the majority and minority of the population. Different lines represent biological replicates. (**A-B**) Frequency dynamics of *S* against *S*_*B*_ and against *L*. (**C-D**) Total cell count dynamics, separated by each strain in the cocultures. (**E-F**) Growth rates over time for each strain in the cocultures, calculated as the log-slope between adjacent timepoints, using the second timepoint as the x-axis location. Insets represent growth rates in stationary phase, from around 8 to 24 hours. Error bars represent standard errors.

We initially focus on the dynamics of strains from 6.5k generations (Figure <u>2</u>). Overall, it is immediately clear that there are larger differences in dynamics in the *L*-*S* cocultures compared to the *S*_*B*_-*S* cocultures. When *S* is in both the majority and minority, *L* has a long, two hour lag time, while *S* starts growing almost immediately (Figure <u>2</u>C-D), causing a large upward spike in *S* frequency. When *S* is cocultured with *S*_*B*_, we don’t see any noticeable lag time; however, when *S* is in the minority, *S* “wakes up” more quickly than *S*_*B*_, leading to a small spike in *S* frequency at the beginning of the time course. We see similar patterns in the cocultures from 17k and 40k generations–both *L* and *S*_*B*_ appear to have growth rates very close to 0 at the beginning of the timecourse, but *S* consistently has a larger initial growth rate (Figure S7).

When 6.5k *L* starts growing, it has a significantly larger exponential growth rate than *S*, pushing the frequency of *S* back down. The magnitude of this growth rate difference is similar regardless of the relative frequency of the ecotypes (Figure 2E-F). In contrast, the differences between *S*_*B*_ and *S* are much smaller. At both starting frequencies, *S*_*B*_ may have a small growth rate advantage compared to *S* early in exponential phase, then *S* appears to grow faster in late exponential phase.

In contrast to the dynamics in lag and exponential phase, the stationary phase dynamics are highly dependent on which ecotype is in the majority. When *S* is cocultured with *L, S* grows better than *L* under both conditions, but the absolute growth rates differ between the conditions (Figure 2E-F). When *S* is in the minority with *L*, both *S* and *L* have net positive growth in stationary phase, although it is higher for *S*, potentially pointing to the favorable conditions of *L*-dominated stationary phase and the putatively large amount of excreted acetate available for exploitation. In contrast, when *S* is in the majority with *L, S* has a smaller, albeit still positive, net growth rate, while *L* has a net negative growth rate in stationary phase. Concordantly, these patterns suggest that *S*-dominated stationary phase is much less hospitable to both *S* and *L*.

We see different stationary phase patterns when *S*_*B*_ and *S* are in coculture, where *S*_*B*_ now performs consistently better than *S* (Figure 2E-F). When *S*_*B*_ is in the majority with *S, S*_*B*_ has a moderately positive net growth rate, while *S* has essentially a net 0 growth rate in stationary phase. Then when *S*_*B*_ is in the minority, both *S*_*B*_ and *S* have net negative growth rates, but *S* declines more than *S*_*B*_. If *S*_*B*_ were more similar to *L*, i.e. an exponential phase specialist that secretes a substantial amount of acetate, we would have expected that *S*_*B*_-*S* and *L*-*S* cocultures would have similar behavior in stationary phase. Instead, *S*_*B*_ appears to have enhanced survival in stationary phase, and decreases the survival prospects of *S*, perhaps because of the reduced availability of acetate. Thus, while *S*_*B*_ doesn’t have a significant advantage over *S* in exponential phase, like *L* has, it compensates with a clear advantage over *S* in stationary phase, essential for coexistence of *S*_*B*_ with *S*.

The results show differences in stationary phase behavior across generations, as well as several conserved features (Figure S7). Similar to the 6.5k strains, when 17k *S* are in the minority with *L, S* has a large positive growth rate during stationary phase, while *L* does not grow. However, when *S* is in the majority with *L*, it’s growth rate is comparable to that of *L*. The 40k *S* and *L* strains show different patterns, where *L* actually generally has a higher stationary phase growth rate. However, this appears to be offset by a large growth advantage of *S* right at the end of exponential phase/beginning of stationary phase; this growth advantage is much larger when *S* is in the minority compared to when it is in the majority. This indicates that the growth advantage of 40k *S* has shifted earlier, potentially because it has adapted to consumed the acetate secreted by *L* much more quickly.

Again, the stationary phase behavior when 17k and 40k *S* and *S*_*B*_ are grown in coculture is noticeably distinct from the behavior of *L*-*S* coculture. Similarly, 17k *S* also does not grow well in *S*_*B*_-dominated stationary phase. And 17k *S*_*B*_ actually has a large positive stationary phase growth rate when *S* is setting the environment, suggesting that *S*_*B*_ has more to gain from stationary phase when it is in the minority compared to vice versa. The picture shifts again with the 40k strains–*S*_*B*_ benefits very little from being in stationary phase, but in contrast, *S* grows well in stationary phase, especially when dominated by *S*_*B*_. This is quite different from the behavior of 40k *S*-*L* cocultures, albeit in a different direction than the strains from the earlier generations. Thus, in 40k cultures, it appears that *S*_*B*_-*S* cocultures act more like *L*-*S* cocultures from earlier generations, where *S*_*B*_ is the clear exponential phase specialist and *S* is the stationary phase specialist.

Together, these results show that growth traits of *L*-*S* cocultures change over evolutionary time, and *S*_*B*_-*S* cocultures are similar in important ways (e.g. lag responses), but also show departures from the original community (e.g. stationary phase behavior) that reveal how the ecological dynamics have shifted with the new, rediversified ecotype.

### Growth traits in novel environments

While we have shown that *S*_*B*_ has distinct growth traits when in coculture with *S* in the evolutionary condition, does *S*_*B*_ also behave differently compared to *S* in novel environments that neither have been in contact with before? If *S* and *S*_*B*_ mostly behave similarly in novel environments, then perhaps the underlying change between the two morphs is targeted only towards traits relevant to the mechanism of ecological differentiation. On the other hand, if such “pleiotropic” effects are widespread, then the underlying metabolic/physiological shift in *S*_*B*_ may be much broader.

To this end, we competed *S*_*B*_ clones against *S* clones for each LTEE timepoint in the same minimal media base as the evolutionary condition (DM), supplemented with different carbon sources (Figure 3). For comparison, we also competed *S* against *L* clones for each LTEE timepoint in each of the conditions. We chose four different carbon sources that support growth of *S, S*_*B*_, and *L* clones from all timepoints and that enter into central metabolism at different points^40^, potentially allowing us to gain insight into global changes in physiology and metabolism. After growing cocultures together for two days in DM25, we diluted them 1:100 in each different media. We kept the cultures in exponential phase, and took two ecotype frequency measurements via flow cytometry: one right before transfer into the new media, and one at the end of exponential phase. As usual, relative fitness was computed as the change in logit frequency.

**Figure 3.**
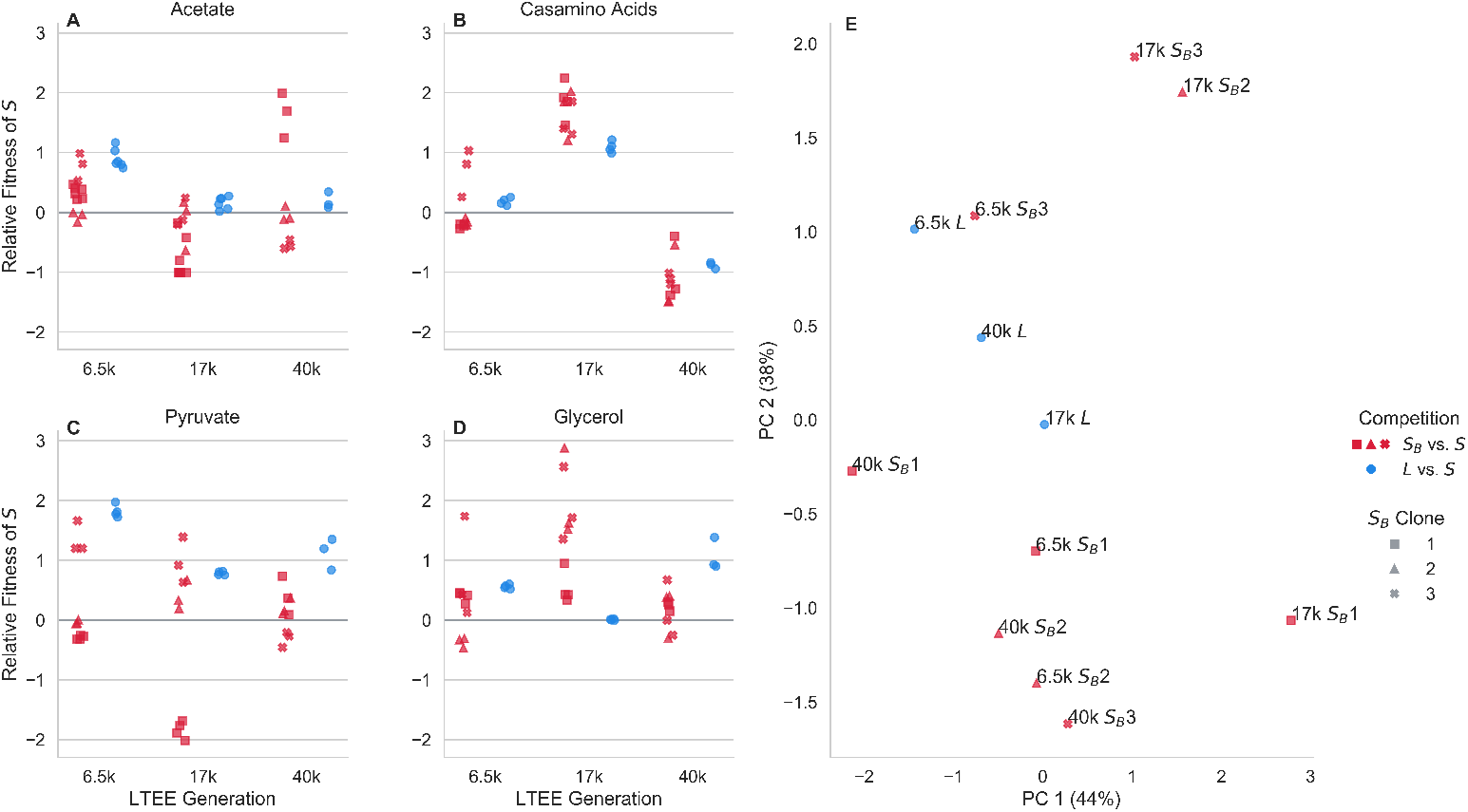
Competition of *S*_*B*_ and *L* against *S* in novel environments. (**A-D**) Red and blue points represent the relative fitness of *S* in competition with *S*_*B*_ and *L* clones from the same LTEE timepoint, respectively, where different symbols represent different clones. Competitions performed in exponential phase in the same media base (DM) supplemented with different carbon sources: (**A**) 200mg/L acetate, (**B**) 1mg/mL casamino acids, (**C**) 20mM pyruvate, (**D**) 20mM glycerol.(**E**) Principal components analysis, using relative fitness in each environment as features. Percentages in parentheses represent percent variance explained by each principal component.

We see that for most *S*_*B*_ clones, across most conditions, *S*_*B*_ is noticeably non-neutral relative to *S*. Perhaps unsurprisingly, and consistent with previous experiments^41^, we see that *L* is also usually non-neutral relative to *S* across the different carbon sources. The relative fitness of *S*_*B*_ and *L* clones varies considerably across timepoints and carbon sources. In general, it appears that there is little relationship between the relative fitness of *S*_*B*_ and that of *L* from the same timepoint. This is also visible in the PCA representation of the data (Figure 3E)–*S*_*B*_ clones within timepoints generally cluster together (but not completely), not with the *L* clones from their timepoint.

Across all three timepoints, we see that *S* is better at growth in acetate compared with *L*. This is consistent with prior findings, and the notion that *S* represents a consistent acetate-scavenging specialist over evolutionary time. In contrast, the behavior of *S*_*B*_ in acetate is more variable, both across time points and between different *S*_*B*_ clones. Most 6.5k *S*_*B*_ clones have a fitness disadvantage in acetate relative to *S* (albeit less pronounced compared to *L*), whereas some 17k and 40k *S*_*B*_ clones have a fitness *advantage* in acetate. This is another sign that *S*_*B*_ is occupying a genuinely different ecological niche compared to *L*.

Again, there is some variation between different *S*_*B*_ clones. For example, the 17k *S*_*B*_ 1 clone behaves noticeably differently compared to the 17k *S*_*B*_ 2 and *S*_*B*_ 3 clones especially in the pyruvate and glycerol conditions, while the three clones cluster together in the acetate condition. The 17k *S*_*B*_ 2 and *S*_*B*_ 3 clones also cluster together, away from the 17k *S*_*B*_ 1 clone in the PCA plot (Figure 3E). The 17k *S*_*B*_ 1 clone also behaved differently compared to the other two in the reciprocal invasion experiment against 17k *L*, where 17k *S*_*B*_ 1 did not show noticeable frequency-dependence (Figure 1D). The 6.5k *S*_*B*_ 3 and 40k *S*_*B*_ 1 clones also cluster away from the other two clones within their timepoint. The conditions where these “outlier” clones diverge from the other clones varies between timepoints–6.5k *S*_*B*_ 3 is different when grown in in pyruvate and casamino acids, and 40k *S*_*B*_ 1 is primarily different in the acetate condition.

Together, these results suggest that the metabolic/physiological change(s) that occur when *S*_*B*_ arises from *S* are not just targeted towards traits relevant for ecological differentiation; rather, there may be global changes to metabolism.

### Transcriptional differences between ecotypes

Given the strong heritability of the *S*_*B*_ phenotype, and multiple traits that differ with respect to *S*, we reasoned that the *S*_*B*_ phenotype may have an underlying genetic cause. Thus, we performed whole-genome shotgun sequencing of several *S* and *S*_*B*_ clones with both short-read sequencing (Illumina) and long-read sequencing (Nanopore) (see Methods). After reference-based assembly, we saw that all *S*_*B*_ clones had several mutations relative to their ancestor, and all *S* clones from the same LTEE generation also has several mutations relative to each other. The mutations were a mix of synoynmous and non-synonymous point mutations, insertions and deletions, and several large genomic rearrangements (see SI section 4). However, none of the mutations differentiated *S* and *S*_*B*_–there were no consistent mutations in specific genes or operons. The large number of mutations separating *S*_*B*_ clones from their *S* ancestor is not surprising; the ara-2 lineage fixed a hypermutator allele before the *S* and *L* lineages split, such that the germline mutation rate is about 100x higher than that of the LTEE ancestor^42^. This makes it likely that many of the mutations are likely (nearly) neutral hitchhikers, or otherwise were not affected by selection. Thus, because of the combination of the high mutational background and lack of detectable genetic parallelism, we cannot determine if the *S*_*B*_ phenotype has a genetic cause, or what the causative mutation(s) would be. If the *S*_*B*_ phenotype is caused by some genetic change, it is likely that many different mutations cause the same/similar phenotype.

To further understand the underlying causes of the *S*_*B*_ phenotype, we turned to measuring transcriptional differences between *L, S*, and *S*_*B*_ from 6.5k generations using RNA-Seq. We chose to focus on 6.5k strains because this is the LTEE timepoint immediately after the *S* and *L* lineages diversified, allowing us to focus on the “minimal” differences between *S* and *L*, rather than after extensive evolution and divergence. We cultured two biological replicates of two independent clones of each *L, S*, and *S*_*B*_ from 6.5k generations in glucose minimal media, and collected samples in mid-exponential phase (see Methods). Procedures for RNA extraction, sequencing, and processing are described in Methods. For a broad overview of the data, we first performed a principal components analysis, using (normalized, transformed) expression for each gene as the features (Figure 4A). We see that the first principal component already captures more than half of the variance between samples, which primarily serves to separate the *L* clones from the *S* and *S*_*B*_ clones. The *S* clones appear to cluster together strongly, with the *S*_*B*_ clones flanking them. Hierarchical clustering also reveals that the *S* clones cluster together, with the *S*_*B*_ 2 clone as the next most similar, and the *S*_*B*_ 1 clone as the outer-most member of the cluster (Figure S8). This suggests that there are more differences between *S* and *S*_*B*_ than there are between the two *S* clones, but there are stronger differences comparing both *S* and *S*_*B*_ with the *L* clones. The same picture emerges if we look at the distribution of log_2_ fold expression changes between different ecotypes (Figures 4B, S9). Comparing *S* and *S*_*B*_ with *L*, there are many genes with a large range of expression changes, both increasing and decreasing in expression. In contrast, there are generally smaller differences between the two *S*_*B*_ clones and *S*. Again, there are larger and more differences between *S*_*B*_ 1 and *S*, compared with *S*_*B*_ 2 and *S*, suggesting some amount of variability between the two *S*_*B*_ clones.

**Figure 4.**
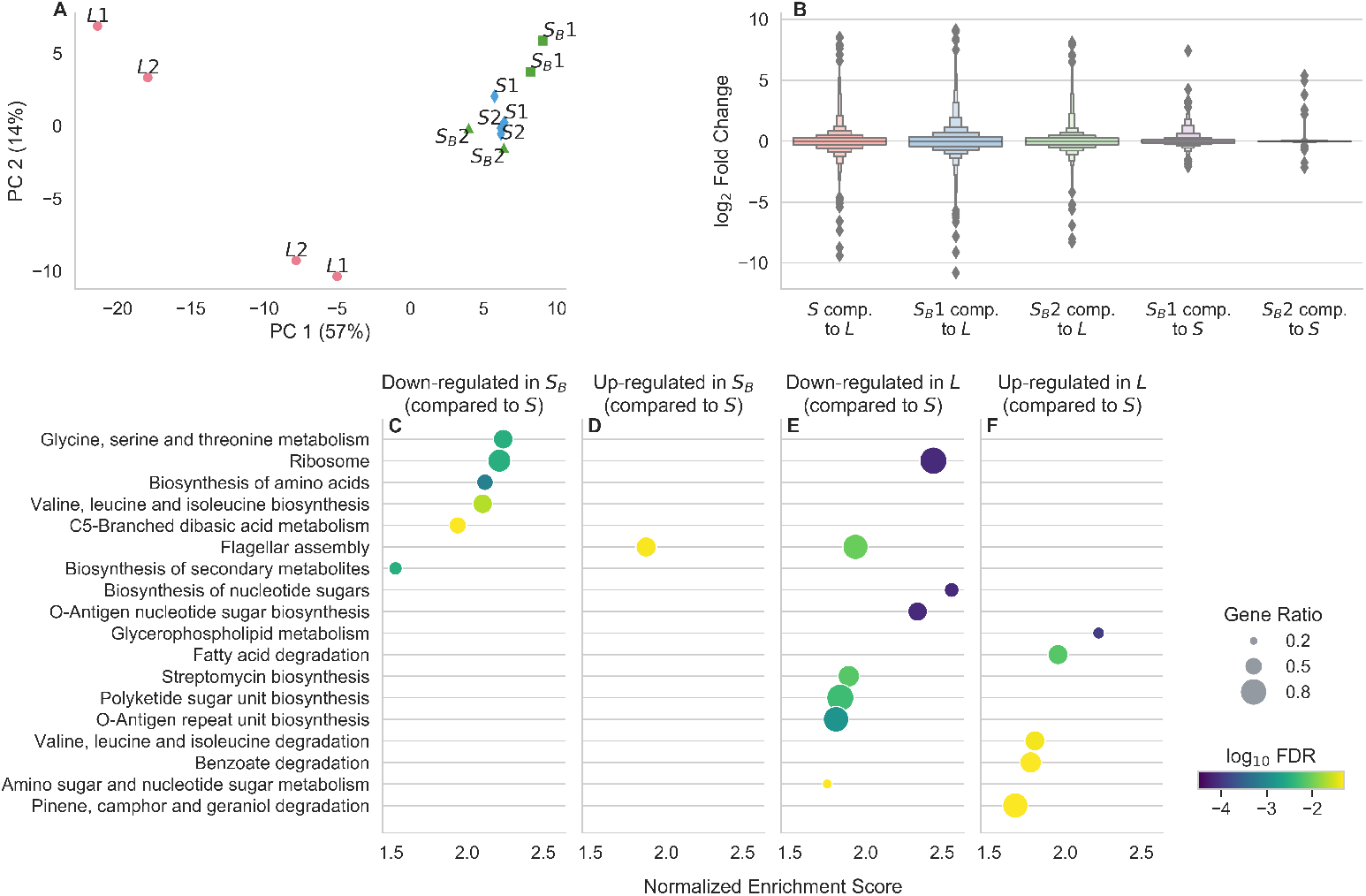
Results from RNA-Seq of *L, S*, and *S*_*B*_ clones from 6.5k generations. (**A**) Principal components analysis of RNA-Seq data, after processing. Samples with the same name represent biological replicates of the same clone; the 1 and 2 labels are to indicate which clone the samples come from. (**B**) Distributions of log_2_ fold changes in gene expression across all genes, comparing different strains to each other. (**C-F**) Results of a KEGG gene set enrichment analysis to identify pathways with coordinated changes in gene expression between ecotypes, where (**C-D**) is comparing *S*_*B*_ to *S* and (**E-F**) is comparing *L* to *S*. Only pathways that are called as significant at *p <* 0.05 after an FDR correction are included; points are colored by FDR-corrected log_10_ p-value. Pathways are ordered by normalized enrichment score, which is roughly a measure of the extent to which pathway-associated genes are overrepresented at the top or bottom of the entire list of genes, ranked by fold expression change. The size of the points is proportional to the “gene ratio”, which is the ratio of core enrichment genes to the total number of genes in the pathway, i.e. the fraction of genes in the pathway that show differential expression.

Given that there are noticeable differences between *S*_*B*_ and *S*, we next sought to understand what those differences represent. Are there identifiable pathways with coordinated expression changes? How do they compare with the differences between *L* and *S*? To this end, we performed gene set enrichment analyses to identify differentially expressed KEGG (Kyoto Encyclopedia of Genes and Genomes) pathways^43^. We first compared *S*_*B*_ to *S* and *L* to *S*, and only look at pathways that are significantly enriched at *p <* 0.05 after a multiple-testing correction (Figure 4C-F). We see that there are a number of pathways significantly down-regulated in *S*_*B*_ compared to *S*, and only one pathway significantly up-regulated (Figure 4C-D). Most of the down-regulated pathways are related to different aspects of amino acid metabolism. We also separately compared *S*_*B*_ 1 and *S*_*B*_ 2 against *S* to better understand the variability between the two clones (Figure S10). As expected, most of the terms identified in the pooled analysis (e.g. ribosomal proteins, amino acid metabolism terms) appeared as the top terms when we analyzed the clones separately, albeit at a lower significance level than when the data from the two clones are pooled together. There are potentially a handful of differences in enriched pathways between the two clones. For example, terms related with O-antigen biosynthesis (e.g. Biosynthesis of nucleotide sugars, O-Antigen nucleotide sugar biosynthesis) may be upregulated in *S*_*B*_ 1, but not *S*_*B*_ 2. The differentially expressed pathways between *L* and *S* are mostly different, there are no terms related to amino acid biosynthesis, and many terms related to lipid metabolism and O-antigen biosynthesis (Figure 4E-F). Differentially expressed pathways in *L* tend to not be differentially expressed in *S*_*B*_, and vice versa (Figure S11).

There are two pathways that are enriched in both comparisons: flagellar assembly and ribosomal proteins. The changes to flagellar assembly expression is in the opposite direction for *S*_*B*_ and *L*, where it is up-regulated in *S*_*B*_ but down-regulated in *L*, suggesting that gene expression for this pathway is ordered *L* < *S* < *S*_*B*_. In contrast, expression of ribosomal proteins is down-regulated in both *L* and *S*_*B*_, perhaps indicating some degree of parallelism involving a fundamental aspect of cell physiology between the two ecotypes. But overall, with the exception of the down-regulation of ribosomal proteins, it appears that the transcriptional changes that differentiate *S*_*B*_ and *L* from *S* are quite distinct.

## Discussion

Our study explores the capacity of an evolved microbial community to quickly regenerate ecological diversity following the removal of a species. Our results suggest that even in the case of a community composed of only two strains in a minimal environment, evolution can leave room for alternative diversification processes.

The rediversified ecotype, *S*_*B*_, demonstrates the robustness of microbial communities to perturbations by sharing several growth traits with the ecotype it replaces, *L*. For instance, both *S*_*B*_ and *L* exhibit slower initial growth or longer lag times compared to *S* across all LTEE timepoints, which may be involved in a trade-off allowing for higher exponential growth rates, as observed in other systems^44^. However, differences between the rediversified and original communities suggest that the mechanism of ecotype coexistence has shifted. Notably, we observe variations in stationary phase responses and survival, as well as distinct patterns of gene expression. Together, these findings indicate that ecological rediversification in the *S*-*L* system may be influenced by a combination of constraints and opportunities. While adaptation may lead some traits to evolve nearly deterministically due to strong ecological or physiological constraints, other trait values may experience more freedom. The interplay between contingency and determinism mirrors patterns observed in various other evolving systems, including the LTEE^45–47^. Dissecting why some traits are more evolutionarily constrained during diversification compared to others could be a fruitful avenue for future investigation.

We attempted to determine a potential genetic origin of the *S*_*B*_ phenotype. However, we did not find any consistent mutations shared between the independent *S*_*B*_ clones, relative to their *S* ancestor. Thus, the *S*_*B*_ phenotype likely either has a large target size, such that many different mutations can cause the same phenotype^48,49^, or it is caused by a non-genetic heritable change. Despite the fact that we did not find any shared mutations, the transcriptional changes of two *S*_*B*_ clones were targeted to the same handful of pathways, predominantly related to amino acid metabolism. This points to parallelism at least on the transcriptional level, if not on the genetic level. Additionally, while the differentially expressed pathways in *S*_*B*_ and *L* relative to *S* were generally different, we saw decreased expression of ribosomal proteins in both ecotypes. The fraction of the proteome devoted to ribosomes is known to control many growth traits in bacteria^50,51^, so the similar changes in *L* and *S*_*B*_ may help to explain the handful of observed similarities in growth traits. One might expect that ribosome expression should be lower in *S*, due to its slower exponential growth rate^52,53^; so the fact that this is not the case may suggest that *S*_*B*_ and *L* are both allocating their proteome not just to optimize exponential growth rate, but also other growth traits as well.

While we saw that *S* could rediversify following isolation, we did not see any obvious ecological or phenotypic diversification when *L* was isolated. There may be several reasons for this. (i) *S* may have some amount of physiological/ genetic/ metabolic plasticity that allows it to diversify that *L* lacks. (ii) Diversification of *L* may happen slowly or rarely, or more quickly only under certain environmental conditions. (iii) Perhaps *L* can rapidly diversify, but cryptically, where no phenotypic changes are obvious without more extensive phenotyping. It is certainly the case that we would not have found *S*_*B*_ without the obvious changes in colony size. It could be that rediversification is much more common than currently appreciated, but simply not detected. Sequencing technologies, including metagenomic^35^ and DNA barcoding-based methods^54^, could help to better reveal the full extent of rediversification across microbial communities. In fact, through metagenomic sequencing, we now know that ecological diversification is much more common in the LTEE than previously thought^35^.

Our study has implications for our understanding of the ecological consequences of species removal or extinction. The ability of microbial communities to rediversify following such perturbations may represent a crucial mechanism by which communities can maintain their functioning and stability over time. While one might think that evolution would be too slow compared to ecological processes, we see here that evolution is crucial for community recovery following a perturbation. The presence of alternative evolutionary pathways, even in a maximally reduced community of only two strains, suggests that such mechanisms may be even more pronounced in communities with greater species richness. In conclusion, our study provides insights into the capacity of microbial communities to regenerate ecological diversity and adapt to environmental perturbations. Further research into the mechanisms that govern these dynamics will be crucial for understanding the functioning and stability of microbial communities, as well as their response to environmental change.

## Methods

### Growth Conditions and Media

Most of the experiments presented here were performed in Davis Minimal Media (DM) base [5.36 g/L potassium phosphate (dibasic), 2g/L potassium phosphate (monobasic), 1g/L ammonium sulfate, 0.5g/L sodium citrate, 0.01% Magnesium sulfate, 0.0002% Thiamine HCl]. The media used in the LTEE and the competitions shown in Figures 1 and 2 is DM25, that is DM supplement with 25mg/L dextrose.

For competition experiments, generally we first innoculated the strain into 1mL LB + 0.2% dextrose + 20mM pyruvate (which we found prevented the emergence of the *S*_*B*_ while allowing for robust growth). After overnight growth, we washed the culture 3 times in DM0 (DM without a carbon source added) by centrifuging it at 2500x*g* for 3 minutes, aspirating the supernatant, and resuspending in DM0. We transferred the washed culture 1:1000 into DM25 in a glass tube. If a strain was isolated directly from a colony, we would instead directly resuspend the colony in DM25. Generally, we grew 1mL cultures in a glass 96 well plate (Thomas Scientific 6977B05). We then grew the culture for 24 hours at 37°C in a shaking incubator. The next day, we transferred all the cultures 1:100 again into 1mL DM25. After another 24 hours of growth under the same conditions, we would mix selected cultures at desired frequencies, then transfer the mixture 1:100 to DM25. After another 24 hours of growth under the same conditions, we would transfer the culture 1:100 to a desired media and start taking flow cytometry measurements–in the competitions of Figures 1 and 2, the media would be DM25, for the competitions of Figure 3, the media would be DM supplement with 200mg/L acetate, 1mg/mL casamino acids, 20mM pyruvate, or 20mM glycerol. For the competitions of Figure 1, we took measurements for 3-4 total days, doing 1:100 serial transfers every 24 hours in DM25; for Figure 2 we took measurements approximately every hour for 8 hours, then another measurement at 24 hours; for Figure 3 we took a second measurement after 8 hours, when the cultures were still in exponential phase.

### Integration of fluorescent proteins

We sought to use flow cytometry to quantify ecotype abundances, which would necessitate that we could differentiate the strains via fluorescence. We decided to integrate fluorescent proteins into a neutral genomic location of our various strains rather than using plasmids, because plasmids can carry a significant metabolic burden, and it is often necessary to add antibiotics to the media to select against plasmid loss. We used a system based on that of Schlechter *et al*.^55^ to integrate fluorescent proteins with miniTn7, a transposon that inserts cargo at a putatively neutral intergenic site downstream of *glmS*. Briefly, the system works by mating the recipient strain-of-interest with a donor strain, harboring a plasmid with the miniTn7 proteins, an ampicillin-resistance gene, a temperature-dependent origin of replication, and the cargo flanked by the left and right Tn7 recognition sites. In this case, the cargo consists of a fluorescent protein, under the control of a broad host-range promoter, and a chloramphenicol resistance gene, for selection of integration.

Out protocol for integration proceeded as follows. First, we grew the donor strain with the desired plasmid in LB + 100*μ*g/mL carbenicillin + 10*μ*g/mL chloramphenicol at 30°C shaken, overnight. We also grew the recipient strain overnight in DM2000 media at 37°C, directly from glycerol stock. The next day, we washed the donor culture by centrifuging it at 2500x*g* for 3 minutes, aspirating the supernatant, and resuspending in DM0. We then measured the optical density (OD) of both cultures, and mixed about 1 OD of each culture on a 20mL LB/agar plate supplemented with 0.2% dextrose + 20mM pyruvate. The cultures were allowed to grow into a lawn overnight at 30°C, allowing the donor strain to conjugate with the recipient. Afterwards, we scraped up the lawn and resuspended it in 3mL DM0. We washed the resuspended culture 3 times, as previously described, and then streaked out the culture on a DM2000 + 10*μ*g/mL chloramphenicol + agar plate, then allowing the plates to incubate overnight at 37°C. This step simultaneously selects against the presence of the donor (the donor is a proline auxotroph), against the Tn7 plasmid (it has a temperature-sensitive origin of replication), and for integration of the Tn7 cargo (via the chloramphenicol resistance gene). After two days of growth, we restreaked a number of colonies that appeared on DM2000/agar plates for isolation. We then tested for integration of the Tn7 cargo by amplifying and sanger sequencing the junction between the genome and the fluorescent protein insertion (see SI section 3 for oligonucleotide sequences), and by looking for fluorescence via fluorescence microscopy. We confirmed that the plasmid was not present in the colony by testing resistance against carbenicillin. We ensured that the colony was not the donor or a contaminant by checking colony morphologies on tetrazolium -maltose (TM), -arabionse (TA), and -xylose (TX) agar plates. We further confirmed identity by sanger sequencing the *arcA* and *aspS* loci of the clones we moved forward with (see SI section 3 for oligonucleotide sequences).

We found that the fluorescence provided by the plasmids designed in Schlechter *et al*.^55^ were insufficiently strong for our purposes. We also needed two different fluorescent proteins with non-overlapping fluorescence profiles so that we could distinguish the two in our flow cytometer. We decided to use the fluorescent proteins sYFP2^56^ and eBFP2^57^ because they share the same ancestor and are highly homologous, and are thus likely to have the same or similar physiological effects on their host, and they have sufficiently different fluorescence profiles that are compatible with our flow cytometer. Thus, we sought to increase the expression levels of the fluorescent proteins, and add in BFP, by constructing new plasmids. We chose to use the strong BBa_J23119 promoter^58^ and a ribosome binding site (RBS) designed *in silico* with the Salis lab “RBS calculator”^59^, placing them immediately upstream of the fluorescent protein sequences. We used gibson assembly to construct the plasmids by ordering compatible oligonucleotides with the promoter and RBS sequences on them, and then using the backbone of pMRE-Tn7-133 from^55^ and the eBFP2 gene from pBad-EBFP2^57^ for the BFP plasmid (see SI for plasmid and oligonucleotide information). Final plasmid sequences were confirmed with sanger sequencing.

### Flow cytometry

For all population measurements taken with flow cytometry, we used the ThermoFisher Attune Flow Cytometer (2017 model) at the UC Berkeley QB3 Cell and Tissue Analysis Facility (CTAF). For every measurement, we loaded the samples into a round bottom 96 well plate, for use with the autosampler. Typically we diluted the samples 1:5 in DM0, but we changed the dilution rate over the course of the 8 hour within-cycle timecourse. We set the flow cytometer to perform one washing and mixing cycle before each measurement, and ran 50*μ*L of bleach through the autosampler in between each measurement to ensure that there was no cross-contamination between wells. We used the “VL1” channel to detect eBFP2 fluorescence, which uses a 405nm laser and a 440/50nm bandpass emmission filter. We used the “BL1” channel to detect sYFP2 fluorescence, which uses a 488nm laser and a 530/30nm bandpass emmission filter. For the triple competitions shown in Figure 1E, we used a BFP-tagged *S*, a YFP-tagged *S*_*B*_, and a non -fluorescent *L* strain. To estimate the frequency of *L*, we added 5 *μ*M of Syto62 red fluorescent dye (ThermoFisher S11344) to the sample immediately before measurement. We used the “RL1” channel to detect Syto62 fluorescence, which uses a 637nm laser and a 670/14nm bandpass emmission filter. We always used a sample flow rate of 25*μ*L/min.

To analyze flow cytometry data, we first create threshold gates to sufficiently separate the “noise cloud” (nonfluorescent particles present even when running blank media) from particles with clear fluorescence. We noticed that in addition to seeing single positive BFP^+^ and YFP^+^ particles, we also see some particles called as fluorescent in both channels (Figure S3). We observed that the proportion of double positive events decreased as a function of fluid flow rate and dilution rate (Figure S4), suggesting that sometimes multiple cells end up in front of the flow cytometry laser at the same time, and are counted as one event. Thus, we sought to correct for this effect. We will have to make an assumption that the probability of a cell ending up in front of the laser is constant per unit time, and uncorrelated in time, i.e. that it is a poisson process. Thus, for any given window of time, the probability of observing some number of events is distributed as a poisson distribution. So under this model, the observed BFP or YFP “clouds” will consist of single cells, double cells, triple cells, and so on. Similarly, there are many combinations of BFP/YFP cells that can end up in the double positive cloud. So in order to get the expectation of the observed frequencies, we add up the contributions of singlets, doublets, triplets, etc by considering the probability of *n* cells passing in front of the laser together times the probability of all *n* cells being the same color,

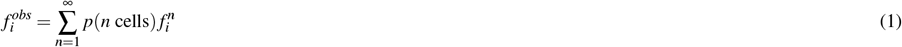

where *i ∈{1,2}*. As previously mentioned, *p* (*n* cells) will follow a poisson distribution, but as we do not observe the case when zero cells pass in front of the laser, we will use a zero-truncated poisson.

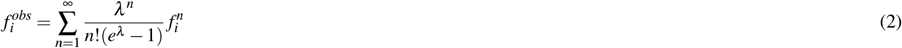

Where *λ* is the average number of cells per event. We have two equations (for 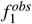 and 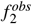) and two unknowns (*λ* and *f*_1_), so we can solve for the real frequencies, which we solve for via numerical root-solving, performing the sum to 100 (which appears to be more than sufficient for convergence). The total cell count *N* also must be corrected, where *N*_*corrected*_ = *N*_*observed*_*λe*^*λ*^ */*(*e*^*λ*^ 1). The post-correction frequencies appear to be well-reflective of frequencies measured with colony counting (Figure S1).

### Whole Genome Sequencing

To perform short-read sequencing of *S*_*B*_ and *S* clones (see SI), we first grew the clones overnight in 1mL of DM2000, then pelleted the cultures and extracted genomic DNA with the DNeasy Blood and Tissue Kit (Qiagen 69504). We prepared the sample libraries with NEBNext DNA Library Prep kit for Illumina according to the manufacturer’s protocol (New England Biolabs E7645). We sequenced the samples with the Illumina 4000 HiSeq 150PE. We used `breseq`^60^ to compare raw reads to the REL606 genome^61^ (GenBank: CP000819.1) and to the *S* ancestor of each *S*_*B*_, and then call genetic variants.

To perform long-read sequencing of *S*_*B*_ and *S* clones (see SI), we again grew the clones overnight in 1mL of DM2000, then pelleted the cultures. High-molecular weight DNA extraction was performed via a standard phenol-chloroform extraction and isopropanol precipitation. Distribution of DNA fragment sizes were obtained using the Agilent Femto Pulse System. Fragment size selection was performed using Pippin Prep (Sage Biosciences). The samples were prepared for sequencing with the Nanopore ligation sequencing kit (Oxford Nanopore, SQK-LSK109). The libraries were then sequenced on an Oxford Nanopore MinION. We used `minimap2`^62^ and `sniffles`^63^ with default parameters to detect structural variants.

### RNA Sequencing

6.5k *S* and *L* clones 1 and 2 were isolated from REL11555 and REL11556 respectively; 6.5k *S*_*B*_ clones 1 and 2 were the same clones as previously described. Cultures of 6.5k *S, S*_*B*_, and *L* clones 1 and 2 were started directly from glycerol stock into 1ml LB + 2g/L dextrose + 20mM pyruvate, as a pre-culture. We started two independent cultures for each clone as biological replicates. After overnight growth, the cultures were washed by centrifuging the cultures at 2500x*g* for 3 minutes, aspirating the supernatant, and resuspending in DM0, repeated three times. Then, the cultures were diluted 1 : 10^−4^ into 1mL fresh DM media supplemented with 4g/L dextrose, in glass tubes. After approximately four hours of growth at 37°C, the cultures were again diluted 1 : 50 in 1mL of the same media in glass tubes. The cultures were grown shaken at 37°C. The cultures were grown to mid-exponential phase, i.e. until *OD∼* 0.4, then the entire culture was immediately centrifuged at 2500x*g* for 3 minutes to pellet. Immediately after centrifugation, we resuspended the pellets in 25*μ*L TES buffer (10 mM Tris-HCl [pH 7.5], 1 mM EDTA, and 100 mM NaCl) and then lysed the pelleted cultures with 250U/*μ*L lysozyme (Ready-Lyse Lysozyme Solution; Lucigen R1804M) at room temperature for 5 minutes. For all subsequent steps, we used Monarch Total RNA Miniprep Kit (New England BioLabs T2010S) according to the standard given protocol for gram-negative bacteria. Samples were eluted in 30*μ*l nuclease-free water, and stored at -80°C. The concentration and purity of all RNA samples was quantified using Qubit.

RNAse-free DNAse (Invitrogen AM2222) was used to treat the samples for DNA removal. The library preparation was conducted using Illumina’s Stranded Total RNA Prep Ligation with Ribo-Zero Plus kit and 10bp IDT for Illumina indices. Subsequently, the samples were sequenced using NextSeq2000, resulting in 2×51bp reads. The process of demultiplexing, quality control, and adapter trimming was carried out using `bcl-convert` (v3.9.3) and `bcl2fast` (v2.20.0.445) (both are proprietary Illumina software for the conversion of bcl files to basecalls). `HISAT2` (v2.2.0)^64^ was used for read mapping. Reads were mapped to the REL606 genome^61^ (GenBank: CP000819.1). The read quantification was performed using the functionality of `featureCounts` (v2.0.1) in `Subread`^65^. All of the above steps in the pipeline were performed with default parameters, the last two steps also were run with `-very-sensitive` and `-Q 20` tags, respectively. All sequencing and pre-processing steps were performed by SeqCenter, LLC.

After pre-processing, we obtained a matrix of read counts for each gene for each sample. With this table, we used `DESeq2` (v1.38.3)^66^ to compute fold change in expression between strains and variance-stabilized relative expression values for each gene across samples (blindly with respect to the design matrix), all with default parameters. We used the variance-stabilized relative expression values for the principal components analysis (PCA). We used the `ashr` method (v2.2)^67^ with default parameters to shrink and regularize the log_2_ fold changes in expression. We computed log_2_ fold change in expression between samples in two ways, (1) treating the *S*_*B*_ clones as one “strain”, and (2) treating the *S*_*B*_ clones as separate, so that we get different fold changes in expression for each clone. Otherwise, for *S* and *L*, we pooled data across the two clones and biological replicates when computing fold change in expression. We used the package `clusterProfiler` (v4.6.2)^68^ to perform the KEGG gene set enrichment analysis (GSEA). We used the previously computed log_2_ fold change in expression as the metric to pre-sort the list of genes. We used the `gseKEGG` method along with the parameters `organism=“ebr”`, `nPerm=1000000`, `minGSSize=3`, `maxGSSize=800`, `eps=1e-20` to perform the analysis. We also generated the GSEA plots (Figures S12,S13) with `clusterProfiler`.

## Acknowledgements

We thank Adam Arkin, Morgan Price, Benjamin Good, Tanush Jagdish, Michael Desai, Jeff Barrick, Dominique Schneider, and all members of the Hallatschek lab (past and present) for helpful comments and advice on the project. We thank Richard Lenski for sending us the LTEE-derived strains and populations, along with experimental advice and feedback. Research reported in this publication was supported by a National Science Foundation CAREER Award (1555330). This work was supported by the National Institute of General Medical Sciences of the NIH under award R01GM115851 and by a Humboldt Professorship of the Alexander von Humboldt Foundation. JAA acknowledges support from an NSF graduate research fellowship, a Berkeley fellowship (from UC Berkeley), and Lloyd and Brodie scholarships (from UC Berkeley Dept of Bioengineering). We thank Mary West of the Cell and Tissue Analysis Facility (CTAF) at UC Berkeley. This work was performed in part in the QB3 CTAF, that provided the ThermoFisher Attune Flow Cytometer (2017 model). RNA sequencing and processing was performed by SeqCenter, LLC. Nanopore library preparation and genomic sequencing along with Illumina sequencing was performed by the Vincent J. Coates Genomics Sequencing Laboratory at UC Berkeley, supported by NIH S10 OD018174 Instrumentation Grant.

