## Supplementary Information for "Rediversification Following Ecotype Isolation Reveals Hidden Adaptive Potential"

---

#### Supplementary Information

Joao A Ascensao, Jonas Denk, Kristen Lok,
QinQin Yu, Kelly M Wetmore, Oskar Hallatschek

2023

### Contents

**1 Table of strains** **2**

**2 Table of plasmids** **3**

**3 Table of oligonucleotides** **4**

**4 Mutations in $S_B$ clones relative to their ancestor** **4**

4.1 6.5k $S_B$ 1 . . . . . . . . . . . . . . . . . . . . . . . . . . . . . 5
4.2 6.5k $S_B$ 2 . . . . . . . . . . . . . . . . . . . . . . . . . . . . . 5
4.3 6.5k $S_B$ 4 . . . . . . . . . . . . . . . . . . . . . . . . . . . . . 5
4.4 6.5k $S_B$ 5 . . . . . . . . . . . . . . . . . . . . . . . . . . . . . 5
4.5 6.5k $S_B$ 6 . . . . . . . . . . . . . . . . . . . . . . . . . . . . . 6
4.6 17k $S_B$ 1 . . . . . . . . . . . . . . . . . . . . . . . . . . . . . 6

### List of Figures

S1 Flow cytometry accuracy . . . . . . . . . . . . . . . . . . . . . . . . . 8
S2 Flow cytometry controls and bias of $S/L$ CFUs . . . . . . . . . . . 9
S3 Example of raw flow cytometry data and gates . . . . . . . . . . . 10
S4 Varying dilution rate and flow rate in the flow cytometer . . . . . 11

S5 Rediversification of $S$ into $S_B$ . . . . . . . . . . . . . . . . . . . . . 12
S6 Relative fitness of $S_B$ in LB . . . . . . . . . . . . . . . . . . . . . . 13
S7 Growth dynamics over 24-hour cycle . . . . . . . . . . . . . . . . . 14
S8 Heatmap and clustering of gene expression patterns . . . . . . . . 15
S9 Volcano plot of differential expression . . . . . . . . . . . . . . . . 16
S10 Kegg gene set enrichment analysis for $S_B$ clones . . . . . . . . . . 16
S11 Comparison of differentially expressed KEGG pathways between $S_B$ and $L$ . . . . . . . . . . . . . . . . . . . . . . . . . . . . . . 17
S12 Gene set enrichment analysis plots, comparing $L$ and $S$ . . . . . 18
S13 Gene set enrichment analysis plots, comparing $S_B$ and $S$ . . . . 19

### 1 Table of strains

| Strain Name | Internal Name | Ancestor | Notes |
|---|---|---|---|
| REL606 | REL606 | | ara- LTEE ancestor |
| 6.5k $S$ 1 | eJA046 | REL11555 | subclone |
| 6.5k $S$ 2 | eJA047 | REL11555 | subclone |
| 6.5k $S$ 3 | eJA048 | REL11555 | subclone |
| 6.5k $L$ 1 | eJA027 | REL11556 | subclone |
| 6.5k $L$ 2 | eJA028 | REL11556 | subclone |
| 6.5k $S_B$ 1 | eJA049 | 6.5k $S$ 1 | Rediversified in DM25 |
| 6.5k $S_B$ 2 | eJA036 | 6.5k $S$ 1 | Sister to 6.5k $S_B$ 4* |
| 6.5k $S_B$ 3 | eJA395 | 6.5k $S$ 1 | |
| 6.5k $S_B$ 4 | eJA037 | 6.5k $S$ 1 | Sister to 6.5k $S_B$ 2* |
| 6.5k $S_B$ 5 | eJA034 | 6.5k $S$ 2 | |
| 6.5k $S_B$ 6 | eJA035 | 6.5k $S$ 3 | Rediversified in DM25 |
| 17k $S$ 1 | eJA052 | REL11557 | |
| 17k $L$ 1 | eJA031 | REL11578 | |
| 17k $S_B$ 1 | eJA055 | 17k $S$ 1 | |
| 17k $S_B$ 2 | eJA397 | 17k $S$ 1 | |
| 17k $S_B$ 3 | eJA398 | 17k $S$ 1 | |
| 40k $S$ 1 | eJA172 | REL10927 | Isolated from mixed population |
| 40k $L$ 1 | eJA174 | REL10927 | Isolated from mixed population |
| 40k $S_B$ 1 | eJA177 | 40k $S$ 1 | |
| 40k $S_B$ 2 | eJA399 | 40k $S$ 1 | |
| 40k $S_B$ 3 | eJA400 | 40k $S$ 1 | |

*6.5k $S_B$ 2 and 4 were isolated from the same rediversification experiment culture–they come from two different colonies from the same timepoint from the same plate.

#### 2 Table of plasmids

| Plasmid Name | Description | Source of parts | Source |
|---|---|---|---|
| pBad-EBFP2 | Plasmid with eBFP2 gene | | [1]; Addgene 14891 |
| pMRE-Tn7-133 | miniTn7 plasmid to insert sYFP2 into genome at attTn7 | | [2]; Addgene 118551 |
| pJA17 | miniTn7 plasmid to insert eBFP2 (with strong promoter) into genome at attTn7 | Backbone from pMRE-Tn7-133; eBFP2 from pBad-EBFP2; BBa_J23119 promoter [3]; RBS designed in silico [4] | This study |
| pJA18 | miniTn7 plasmid to insert sYFP2 (with strong promoter) into genome at attTn7 | Backbone from pMRE-Tn7-133; eBFP2 from pMRE-Tn7-133; BBa_J23119 promoter [3]; RBS designed in silico [4] | This study |

#### 3 Table of oligonucleotides

| Oligo Name | Sequence | Purpose |
|---|---|---|
| ja35 | GAAGAGGAT AAAACCGTGGA | Amplify arcA for genotyping |
| ja36 | AGGTCAGGG ACTTTTGTAC | Amplify arcA for genotyping |
| ja40 | TTCGAAGCG ACAGATGGC | Sanger sequence arcA for genotyping |
| ja37 | CAGTTGTGACATA CAGCTAACGCT | Amplify aspS for genotyping |
| ja38 | GCTCATGGGAGTT CACTCAGTTG | Amplify aspS for genotyping |
| ja42 | ACTCTAACC ACGTCAACACC | Sanger sequence aspS for genotyping |
| ja200 | ATTCGCTAA ACTGTgctagcatta tacctaggactgagct agctgtcaagctgtc cataaaaccgccc | Gibson assembly primer to make pJA17/18 |
| ja201 | atgctagcACAGTTT AGCGAATACGT CATAGAGCATTA AGGAGGTCATAG atggtgagcaagggcgag | Gibson assembly primer to make pJA17/18 |
| ja183 | tccaagctcagctaattacttg tacagctcgtccatgc | Gibson assembly primer to make pJA17 |
| ja184 | cgagctgtacaagtaattagct gagcttggactcc | Gibson assembly primer to make pJA17 |
| ja190 | cttcgccgatcaggatgcg | Downstream of glmS - amplify attTn7 junction with FP |
| ja191 | tcgccctcgaacttcacctc | In pJA17/18 - amplify attTn7 junction with FP |
| ja170 | caaaatcggttacggttgag | Downstream of glmS - sequence attTn7 junction with FP |

#### 4 Mutations in $S_B$ clones relative to their ancestor

All comparison tables generated with (1) `minimap2` [5] and `sniffles` [6] for structural variants, and (2) `breseq` [7] for all other variants.

### 4.1 6.5k $S_B$ 1

| position | mutation | annotation | gene |
|---|---|---|---|
| 242,204 | (G)9→10 | pseudogene (194/373 nt) | ECB_00212 → |
| 942,720 | (C)7→8 | intergenic (+464/+235) | clpA → / ← serW |
| 1,292,775 | A→G | intergenic (-48/-556) | hns ← / → tdk |
| 1,368,422 | (T)8→9 | intergenic (+40/-172) | pspE → / → ycjM |
| 1,444,298 | C→T | R136R (CGC→CGT) | ydbC → |
| 1,829,779 | G→A | intergenic (+11/+82) | pncA → / ← ydjE |
| 2,034,188 | A→G | Y26Y (TAT→TAC) | manC ← |
| 2,165,552 | (C)5→6 | pseudogene (306/1685 nt) | yehU ← |
| 2,704,655 | T→C | F41F (TTT→TTC) | ygaX → |
| 2,771,495 | T→C | T51A (ACC→GCC) | cysN ← |
| 2,998,766 | G→A | intergenic (+40/-66) | yqgA → / → pheV |
| 3,937,360 | A→G | D43G (GAC→GGC) | rffT → |

### 4.2 6.5k $S_B$ 2

| position | mutation | annotation | gene |
|---|---|---|---|
| 752,223 | C→T | intergenic (+723/-124) | ybgG → / → cydA |
| 942,720 | (C)7→8 | intergenic (+464/+235) | clpA → / ← serW |
| 1,186,311 | G→A | L754F (CTT→TTT) | mfd ← |
| 2,195,273 | (C)9→8 | coding (110/837 nt) | yeiG → |
| 2,762,201 | +T | coding (875/1365 nt) | ygbN → |

### 4.3 6.5k $S_B$ 4

| position | mutation | annotation | gene |
|---|---|---|---|
| 752,223 | C→T | intergenic (+723/-124) | ybgG → / → cydA |
| 942,720 | (C)7→8 | intergenic (+464/+235) | clpA → / ← serW |
| 1,186,311 | G→A | L754F (CTT→TTT) | mfd ← |
| 1,995,685 | (C)8→9 | intergenic (+103/+351) | yeeN → / ← asnW |
| 2,004,021 | -1258 bp | 1258 bp deletion | |
| 3,213,279 | A→G | A20A (GCA→GCG) | yhaV → |
| 4,598,479 | T→C | intergenic (+28/+279) | nadR → / ← yjjK |

### 4.4 6.5k $S_B$ 5

| position | mutation | annotation | gene |
|---|---|---|---|
| 242,204 | (G)9→10 | pseudogene (194/373 nt) | ECB_00212 → |
| 1,422,705 | INV | 179,806 bp inversion | |
| 2,567,537 | T→C | L658L (CTA→CTG) | pbpC ← |
| 4,606,996 | C→T | A151V (GCC→GTC) | creC → |

### 4.5 6.5k $S_B$ 6

| position | mutation | annotation | gene |
|---|---|---|---|
| 508,855 | A→G | G32G (GGA→GGG) | glxR → |
| 751,910 | IS1 (–) +8 bp | intergenic (+410/-430) | ybgG → / → cydA |
| 1,038,663 | A→G | V141V (GTA→GTG) | yccR → |
| 1,140,567 | +771 bp | 771 bp insertion | |
| 1,422,704 | INV | 179,807 bp inversion | |
| 1,184,045 | A→G | R312R (CGT→CGC) | ycfS ← |
| 1,480,910 | (A)8→7 | coding (786/1407 nt) | ydcR → |
| 2,178,109 | G→A | S140L (TCG→TTG) | yohF ← |
| 3,726,210 | T→C | intergenic (-70/+10) | cysE ← / ← gpsA |
| 4,288,802 | A→G | intergenic (-24/+35) | alsB ← / ← rpiR |

### 4.6 17k $S_B$ 1

| position | mutation | annotation | gene |
|---|---|---|---|
| 794,722 | (G)10→9 | pseudogene (282/462 nt) | ECB_00735 → |
| 1,292,373 | IS1 (–) +9 bp | coding (346-354/414 nt) | hns ← |
| 1,603,015 | T→C | I90V (ATC→GTC) | ECB_01505 ← |
| 1,680,075 | +770 bp | 770 bp insertion | |
| 1,796,607 | (C)10→9 | intergenic (-53/+55) | celF ← / ← celD |
| 2,032,376 | G→A | T405T (ACC→ACT) | manB ← |
| 2,104,764 | (CCAG)21→20 | pseudogene (238-241/272 nt) | ECB_01992 → |
| 4,447,544 | +34 bp | pseudogene (636/1388 nt) | treB ← |

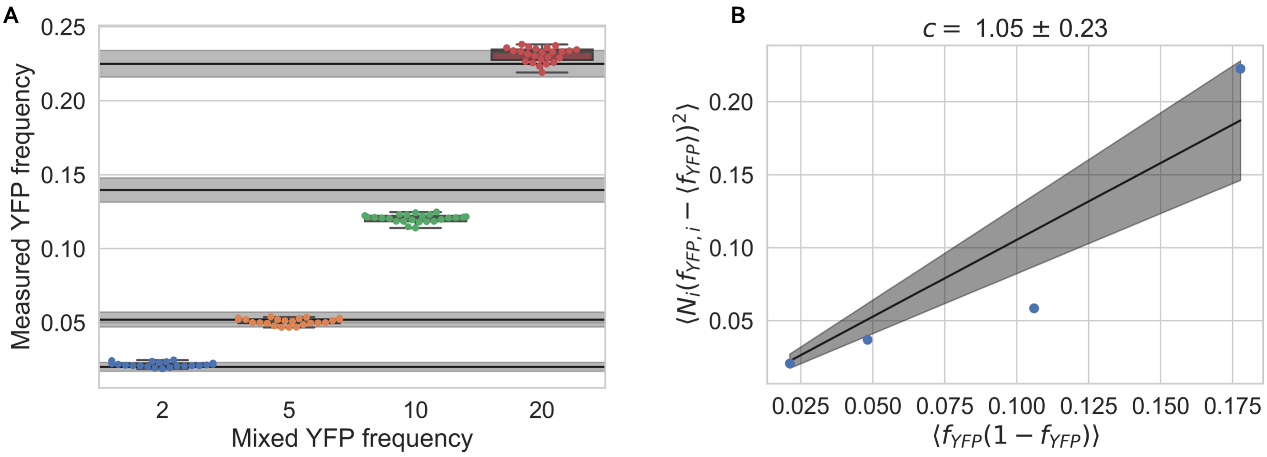


Figure S1: Accuracy of flow cytometer in recovering frequencies of fluorescently-labeled strains in a population. (**A**) We cultured YFP- and BFP-tagged versions of REL606 in DM25, then mixed the cultures at four different frequencies. After culturing for another serial dilution round, we measured each culture 24 times in the flow cytometer (i.e. technical replicates). We also plated the cultures on LB/agar plates and counted the number of blue/yellow colonies for each culture to get CFUs. The colored points represent the flow cytometry measurements. The black lines represent frequencies from CFUs for each of the four cultures, ± standard error. We see that the CFU and flow cytometry measurements mostly match up. (**B**) A plot showing the (scaled) mean-variance relationship, where the variables are transformed such that we would expect a linear relationship with a slope of 1 if the error associated with measuring frequencies followed a binomial, and $\geq 1$ if it was overdispersed. $f_{YFP,i}$ represents the relative frequency of YFP cells in the population, for replicate $i$, and $N_i$ represents the total number of events from the flow cytometer. The blue points are measurements from each experiment, and the line is the fit to the points. Indeed, we see that error looks approximately binomial.

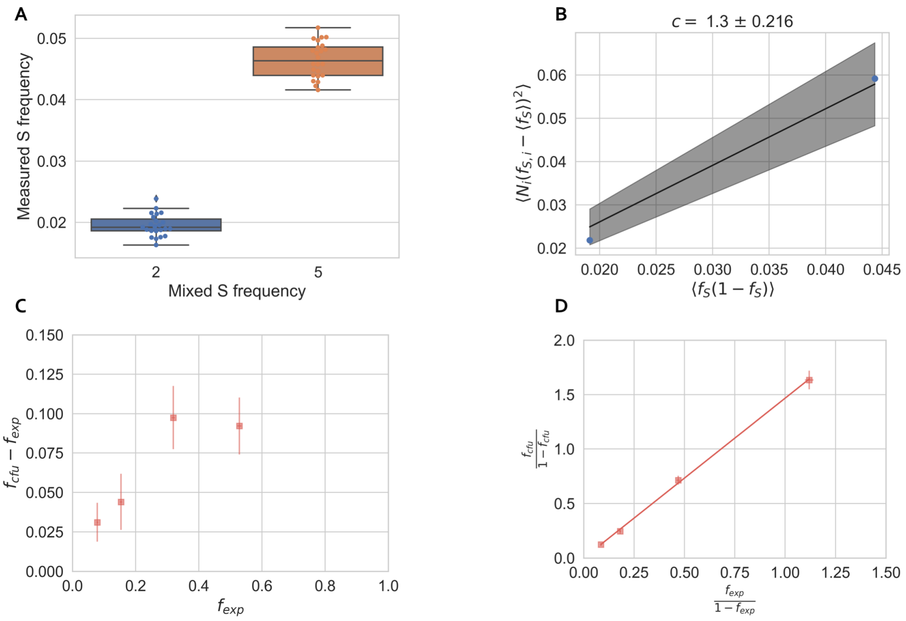


Figure S2: Flow cytometry controls and bias of $S/L$ colony forming units (CFUs). (**A**) We cultured YFP- and BFP-tagged versions of 6.5k $L$ 1 and $S$ 1 respectively in DM25, then mixed the cultures at two different frequencies. After culturing for another serial dilution round, we measured each culture 24 times in the flow cytometer (i.e. technical replicates). The colored points represent the flow cytometry measurements. (**B**) A plot showing the (scaled) mean-variance relationship, similar to Figure S1B. Again, we see that measurement error is consistent with a binomial. (**C**) In a separate experiment, we cultivated four 6.5k $L$ 1 and $S$ 1 cultures at different frequencies, and measured frequencies both by taking a single flow cytometry measurement and by plating the cultures on tetrazolium maltose (TM) plates and counting for CFUs (3 independent plates per culture). We compared the $S$ frequencies from the flow cytometer ($f_{exp}$) to the average $S$ frequencies from CFUs ($f_{cfu}$) and see that $S$ is consistently over-represented in the CFU measurements. (**D**) We hypothesized that there might be a multiplicative bias affecting frequencies. If this was true, then the true frequency would be $f_S = n_S/(n_S + n_L)$ where $n_i$ is the count of ecotype $i$; the biased frequency would be $f_S^{bias} = bn_S/(bn_S + n_L)$ where $b$ is the multiplicative bias. Thus, the odds ratio of the true and biased frequencies should be related linearly, $bf_S/(1 - f_S) = f_S^{bias}/(1 - f_S^{bias})$. We indeed see a linear relationship between the odds ratio of the CFU and flow cytometry frequencies, where $\hat{b} \approx 1.5$. If we trust the flow cytometry measurements more, at least because it more directly measures cell counts and it is unbiased with the LTEE ancestor REL606, then this would imply that measuring $S/L$ frequencies with CFUs incurs a significant multiplicative bias.

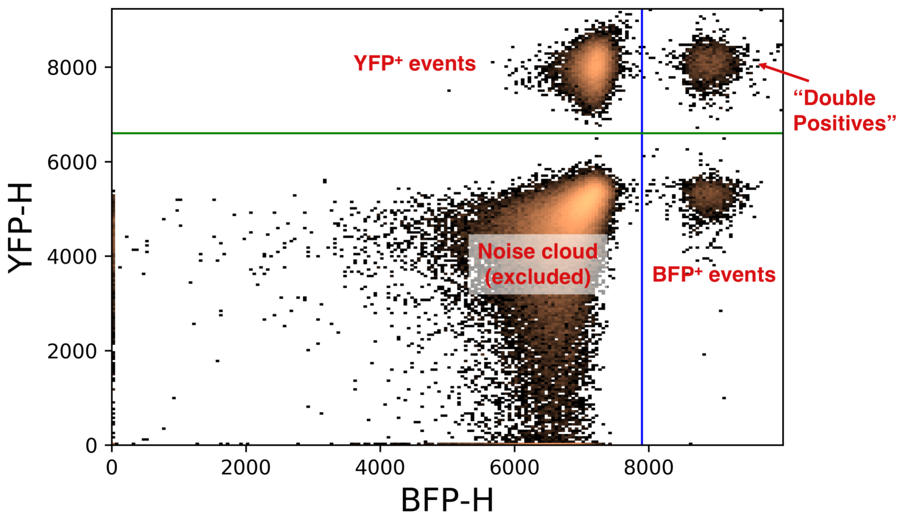


Figure S3: Example of raw flow cytometry data and gates, shown as a 2D histogram of events. The lower left quadrant represents "noise" that is present even when running blank media, and is thus excluded from further analysis. The lower right and upper left quadrants represent events that were called as either BFP or YFP positive, respectively. The upper right quadrant is composed of events that are both BFP and YFP fluorescent; it appears that these events happen when both a BFP and YFP positive cell pass in front of the flow cytometry lasers and are counted as one event. This process appears to be well-described as a poisson process; when we account for this, we appear to recover the correct frequencies (Figure S1). The process by which we account for "double positive" events is detailed in the Methods section.

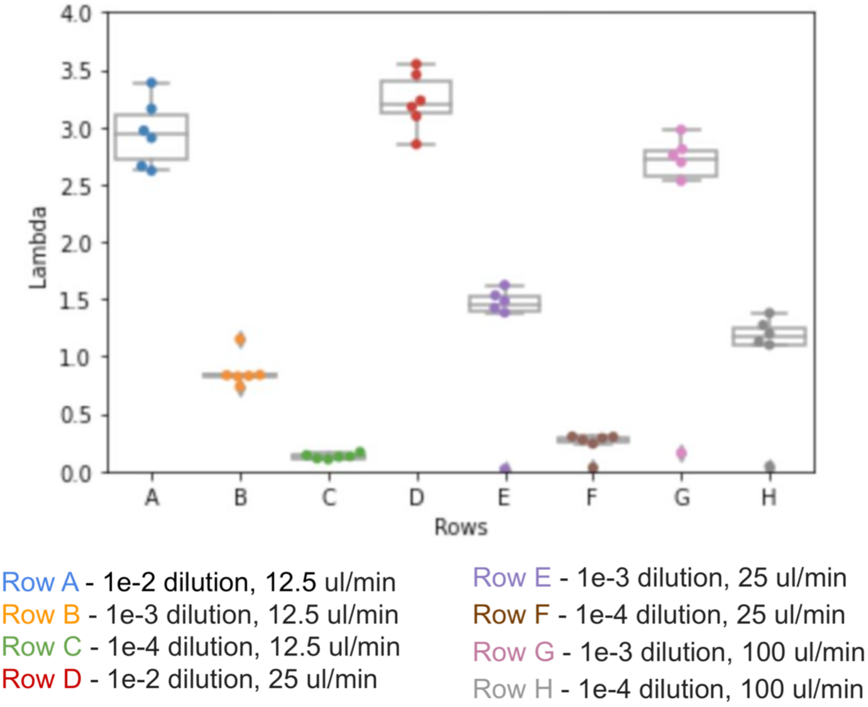


Figure S4: The effect of varying dilution rate and flow rate in the flow cytometer on the average number of cells that end up in front of the laser, i.e. the poisson mean $\lambda$.

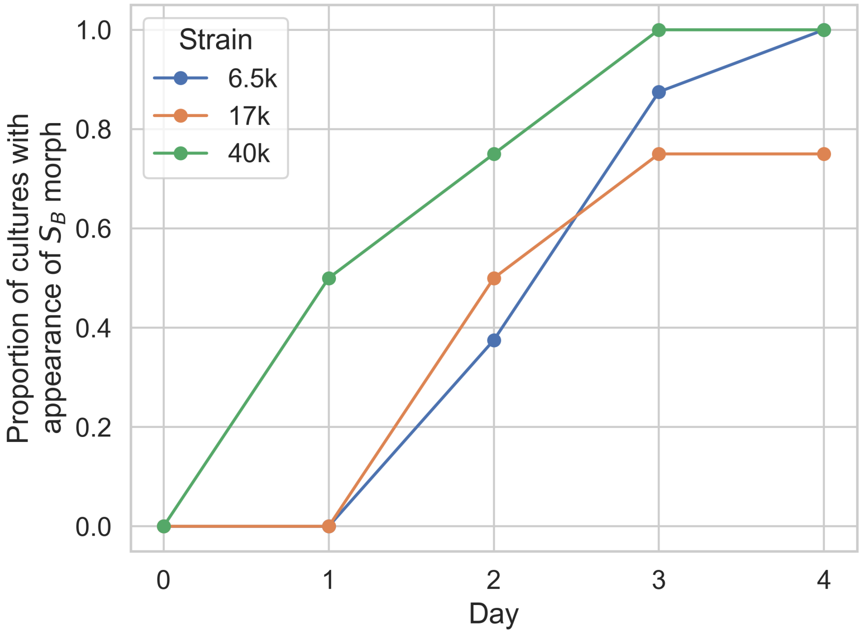


Figure S5: Rediversification of $S$ into $S_B$ in LB. At day 0, we started $S$ clones (picked from DM2000 plates) in 8 replicate 1mL LB cultures, each for $S$ clones from 6.5k, 17k, and 40k LTEE generations. Every day thereafter, we diluted the culture 1:100 into 1mL fresh LB, and we plated the cultures on TA plates (to a final density of around 100 colonies). We then recorded if we saw the appearance of the $S_B$ morph in each of the cultures. We saw the appearance of $S_B$ in most of the cultures across the $S$ clones from the three LTEE timepoints.

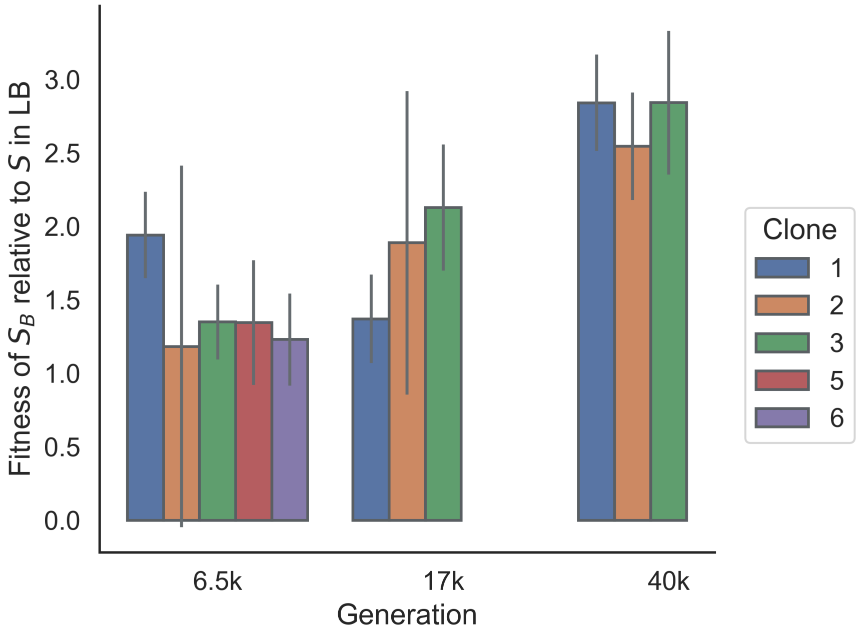


Figure S6: Fitness of $S_B$ clones in LB, relative to $S$. All $S_B$ and $S$ clones were propagated for one day in DM25, then mixed at approximately a 50-50 ratio, and transferred 1:100 to LB. Frequency of $S_B$ was determined by counting colonies on TA plates. Fitness is computed as the change in logit frequency. Error bars represent standard errors.

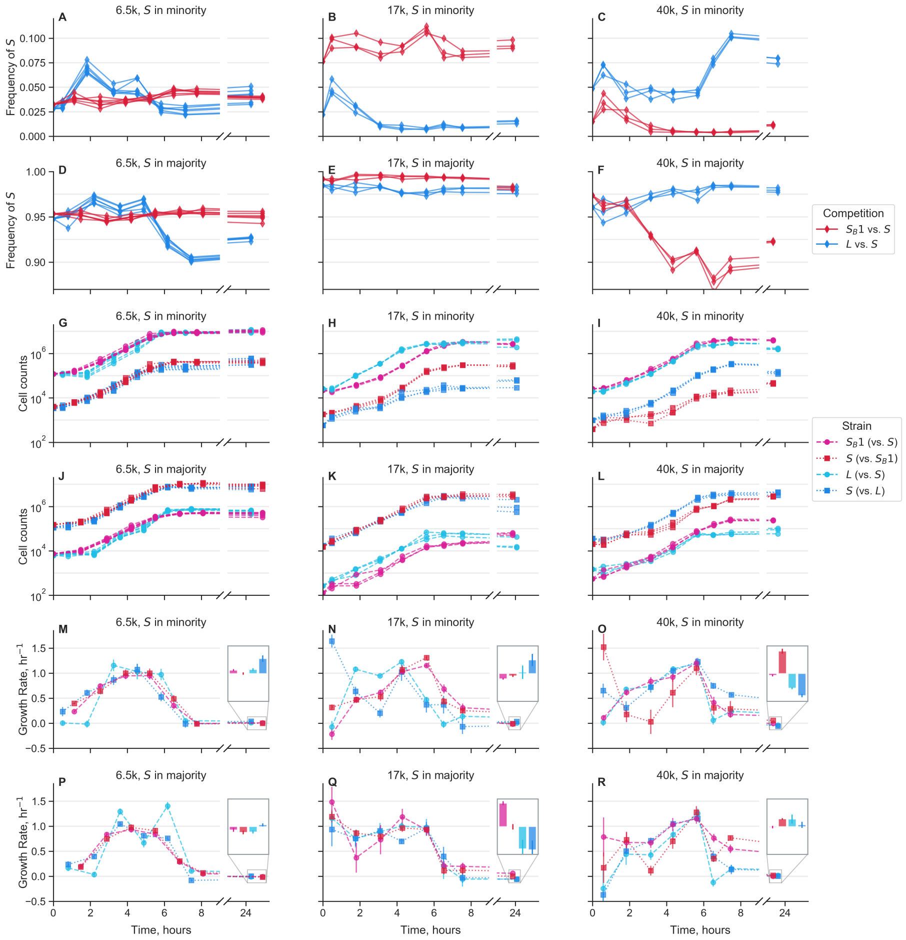


Figure S7: Growth dynamics of cocultures over the course of one twenty-four hour growth cycle. Measurements were taken approximately every hour via flow cytometry for the first eight hours after transfer into new media. An additional measurement was taken approximately 24 hours after the start of the cycle. Mixed $S_B$ with $S$ along with $L$ with $S$ from the same LTEE generation, where ecotypes were mixed both in the majority and minority of the population. All competitions involved $S_B$ used clone 1. Different lines represent biological replicates. Plots in left column represent experiments done with clones from 6.5k generations; 17k clones in the middle column; 40k clones in the right column. (**A-F**) Frequency dynamics of $S$ against $S_B$ and against $L$. (**G-L**) Total cell count dynamics, separated by each strain in the cocultures. (**M-R**) Growth rates over time for each strain in the cocultures, calculated as the log-slope between adjacent timepoints, using the second timepoint as the x-axis location. Insets represent growth rates in stationary phase, from around 8 to 24 hours. All insets use the same y-axis limits, set from -0.13 to 0.13 hr$^{-1}$. Error bars represent standard errors.

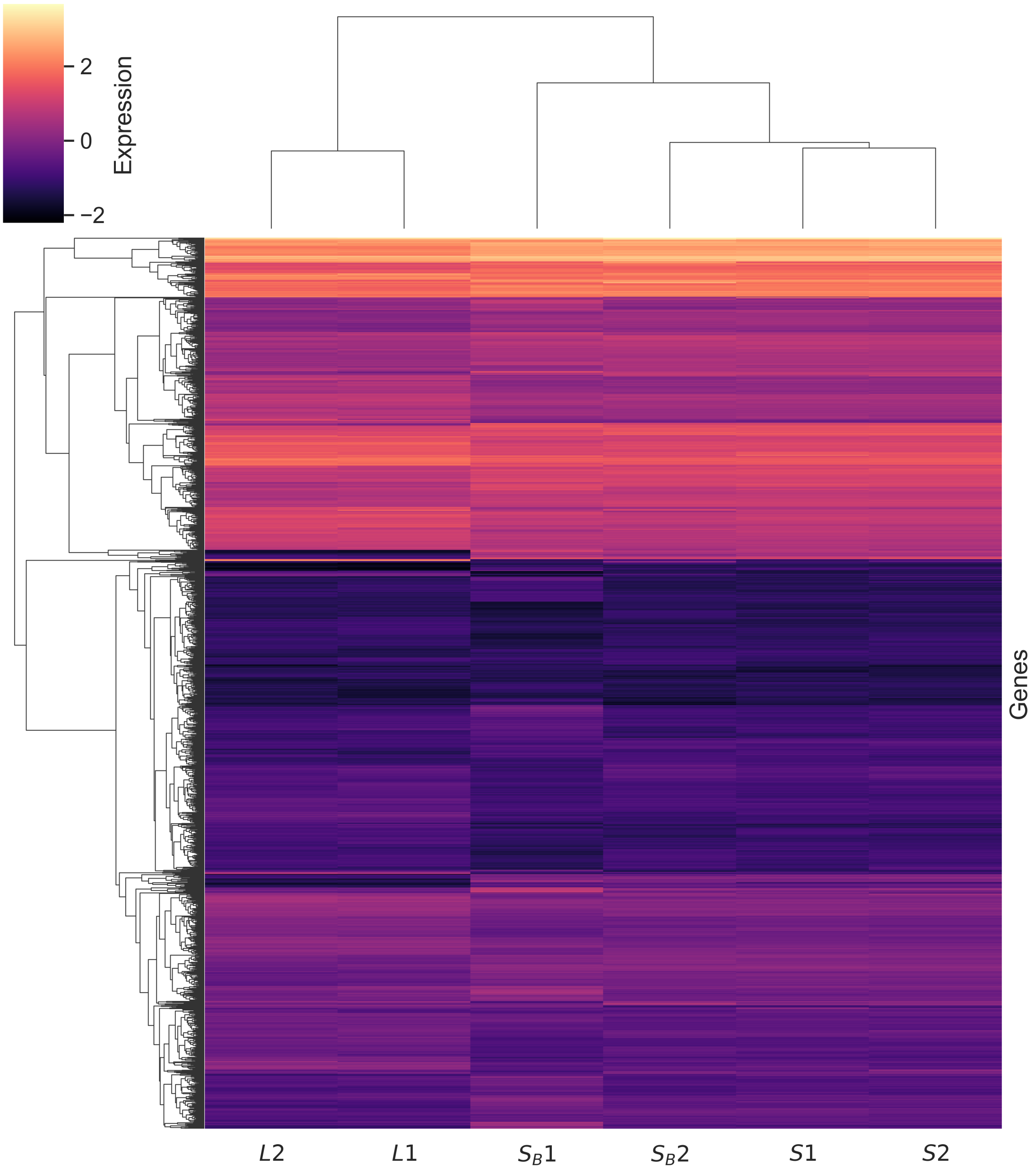


Figure S8: Heatmap and clustering of gene expression patterns. Expression of individual genes is clustered on the y-axis, expression of strains are clustered on the x-axis. We used the variance-stabilized gene expression values from DESeq2, then averaged the two biological replicates for each strain, and centered and scaled the values.

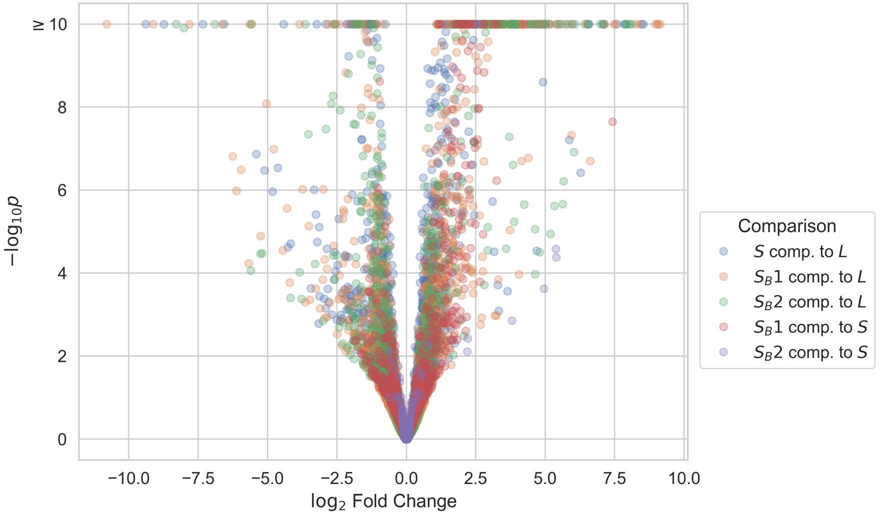


Figure S9: Volcano plot of differential expression, comparing different strains.

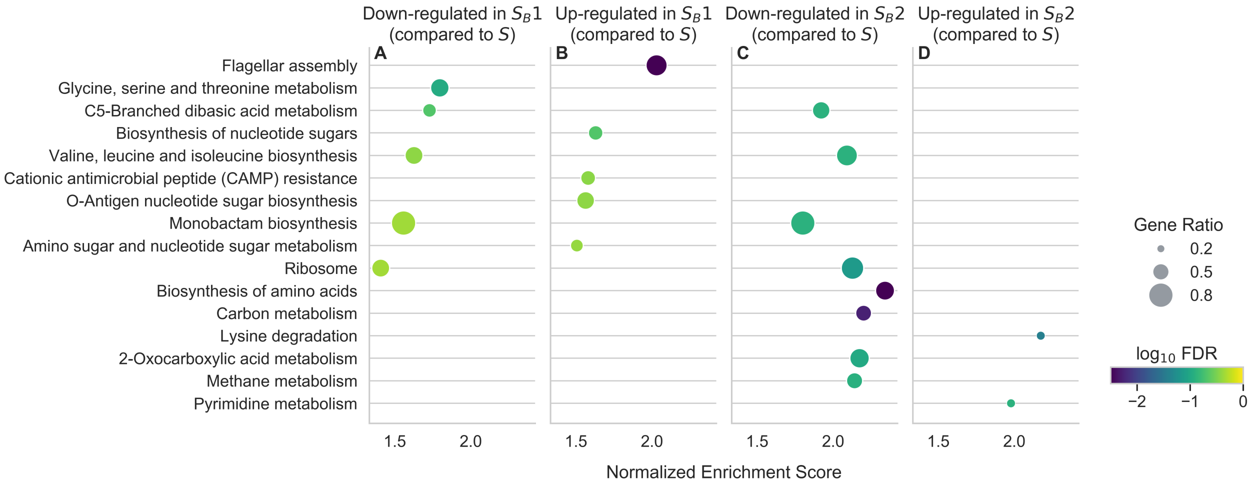


Figure S10: Kegg gene set enrichment analysis for the two $S_B$ clones, showing the top ten genes (sorted by FDR-corrected p-value) for each clone.

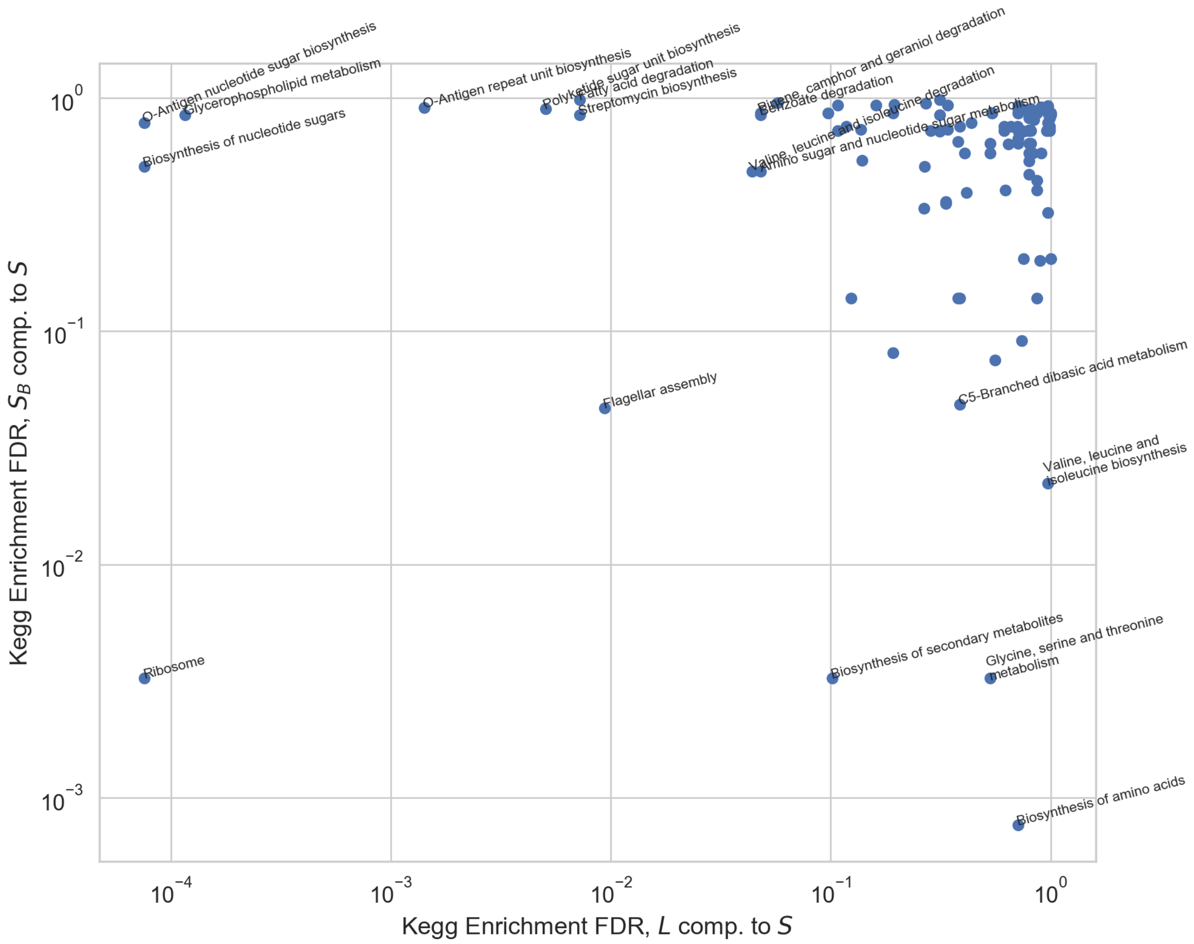


Figure S11: Comparison of differentially expressed KEGG pathways between $S_B$ and $L$. Shown are the FDR-corrected p-values of each KEGG pathway, with the name shown of significantly enriched pathway in either condition. With the exception of the "Ribosome" and "Flagellar Assembly" pathways, the pathways differentially expressed in $S_B$ and $L$ are mostly orthogonal to each other.

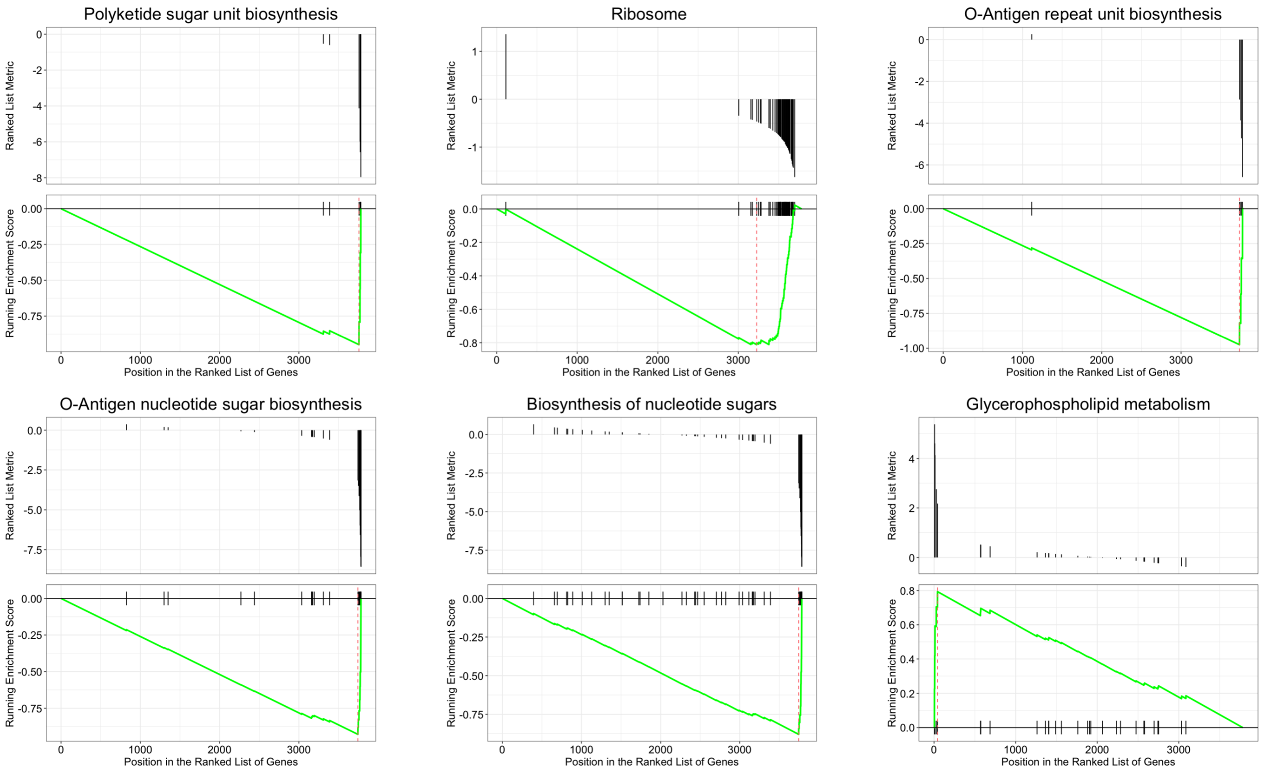


Figure S12: Gene set enrichment analysis plots, comparing $L$ and $S$. Each subpanel represents a different KEGG pathway that was called as significantly enriched when comparing expression patterns of the two strains. The metric used was $\log_2$ fold change in gene expression (from DESeq2) of $L$, compared to $S$.

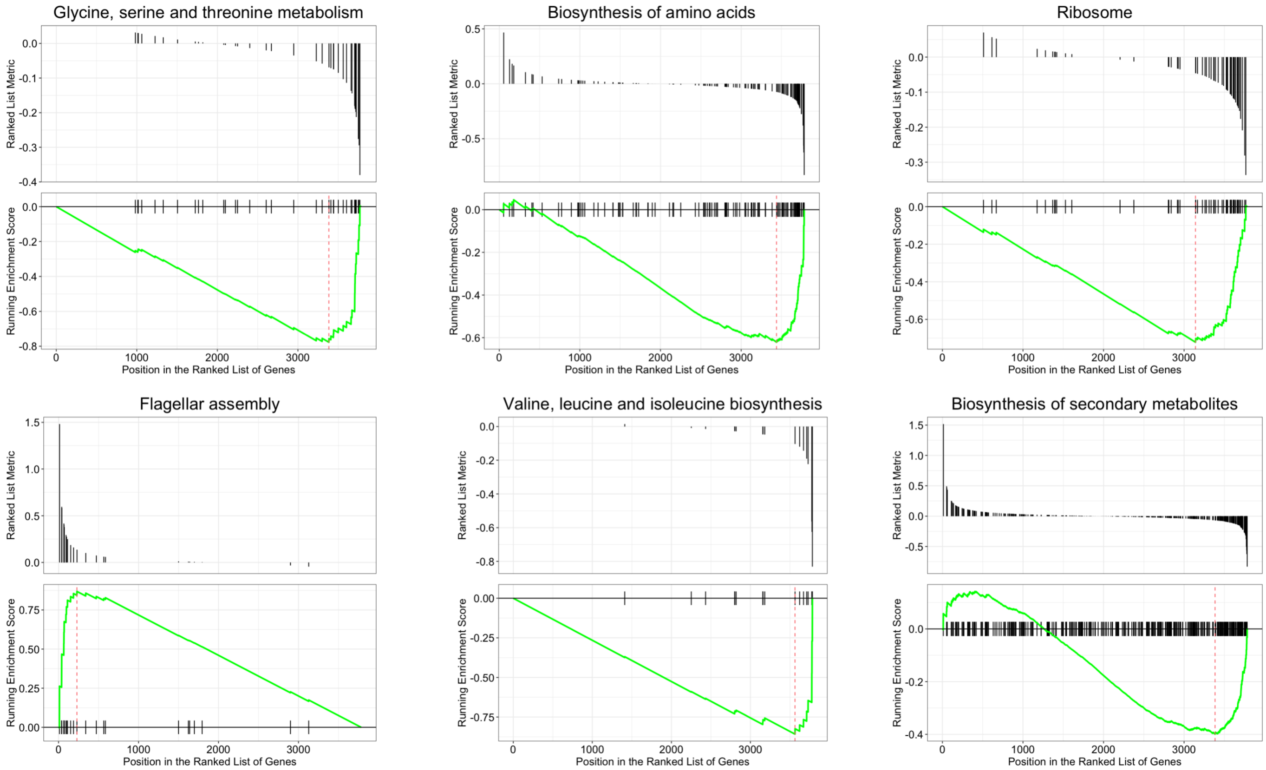


Figure S13: Gene set enrichment analysis plots, comparing $S_B$ and $S$. Each subpanel represents a different KEGG pathway that was called as significantly enriched when comparing expression patterns of the two strains. The metric used was $\log_2$ fold change in gene expression (from DESeq2) of $S_B$, compared to $S$.
